# Integrating brainstem and cortical functional architectures

**DOI:** 10.1101/2023.10.26.564245

**Authors:** Justine Y. Hansen, Simone Cauzzo, Kavita Singh, María Guadalupe García-Gomar, James M. Shine, Marta Bianciardi, Bratislav Misic

## Abstract

The brainstem is a fundamental component of the central nervous system yet it is typically excluded from *in vivo* human brain mapping efforts, precluding a complete understanding of how the brainstem influences cortical function. Here we use high-resolution 7 Tesla fMRI to derive a functional connectome encompassing cortex as well as 58 brainstem nuclei spanning the midbrain, pons and medulla. We identify a compact set of integrative hubs in the brainstem with widespread connectivity with cerebral cortex. Patterns of connectivity between brainstem and cerebral cortex manifest as multiple emergent phenomena including neurophysiological oscillatory rhythms, patterns of cognitive functional specialization, and the unimodal-transmodal functional hierarchy. This persistent alignment between cortical functional topographies and brainstem nuclei is shaped by the spatial arrangement of multiple neurotransmitter receptors and transporters. We replicate all findings using 3 Tesla data from the same participants. Collectively, we find that multiple organizational features of cortical activity can be traced back to the brainstem.

## INTRODUCTION

The brain is a network of functionally interacting neural populations. Studying the functional architecture of the brain in awake humans is possible with multiple imaging technologies, although these technologies are often biased towards the cortex where signal quality is highest [10]. As a result, key findings about functional activity in the brain—including the presence of functionally specialized brain regions [59], networks of regions with synchronized neural activity [109, 127], and mechanisms behind higher-order cognitive processes [3]—are primarily limited to the cerebral cortex. An important question is therefore: what role do extracortical structures have in cortical function?

Perhaps the biggest missing piece of modern *in vivo* brain network reconstruction is the brainstem. This early evolutionary structure is crucial for survival and consciousness, and integrates signals from throughout the nervous system. Furthermore, multiple neurotransmitter systems originate in brainstem nuclei and project throughout the cortex, shaping cortical activity [39, 101, 118]. In stark contrast to research on cortical function, knowledge about brainstem function primarily comes from either lesion studies or studies in model organisms, and these studies are often limited to specific brain-stem nuclei or pathways [37, 74, 76, 81, 82]. Exciting recent imaging advances have improved the feasibility of functional imaging in the whole brainstem including ultrahigh-field magnetic resonance imaging (MRI) scanners and extensive brainstem-specific physiological noise reduction pipelines [10, 17, 27, 103]. Furthermore, recent development of brainstem atlases encompassing multiple nuclei has made it possible to augment the cortical functional connectome with an anatomically comprehensive representation of the brainstem [15, 16, 45, 46, 104, 106].

Here we study how functional activity throughout the brainstem aligns with cortical function by analyzing a high resolution 7 Tesla resting-state fMRI dataset in conjunction with a whole-brainstem atlas spanning 58 nuclei across midbrain, pons and medulla. First we identify hubs of brainstem-cortex connectivity and find that electrophysiological signatures of neural oscillations are reflected by brainstem-cortex functional connectivity. Next, we cluster brainstem nuclei with respect to how they connect to the cortex and identify communities of brainstem nuclei that subserve familiar cortical functional activation patterns related to memory, social cognition, movement and sensation, and emotion. Using PET-estimated brain maps for 18 neurotransmitter receptors and transporters, we find chemoarchitectonic signatures of brainstem-cortex functional connectivity. Finally, we demonstrate that the cortical functional hierarchy delineating unimodal (lower-order) and transmodal (higher-order) brain regions reflects patterns of connectivity with the brainstem. Altogether, using simultaneous *in vivo* human imaging of brainstem and cortical functional activity, this study extends our perspective of cortical function—including dynamics, cognitive function, and the unimodal-transmodal cortical functional gradient—to the brainstem, demonstrating how cortical functional architecture consistently reflects brainstem influence.

## RESULTS

Resting-state functional MRI time-series in the cortex and brainstem were acquired on a 7 Tesla scanner in 20 unrelated healthy participants (29.5 *±* 1.1 years, 10 males, 10 females), and replication data was acquired on a 3 Tesla scanner in the same individuals. Brainstem data were processed following established brainstem-specific protocols, and all functional connections were defined based on specified cortical and brainstem seed and target regions (see *Methods* for details). Cortical regions were defined according to the 400 regions in the Schaefer parcellation [96], and brainstem nuclei were defined according to the 58 nuclei in the Brainstem Navigator atlas (50 bilateral, 8 midline nuclei; Fig. 1a, b; atlas available at https://www.nitrc.org/projects/brainstemnavig) [15, 16, 45, 46, 104, 106]. We first confirm that the nuclei in the brainstem atlas are well aligned with expected receptor densities from positron emission tomography (PET) imaging, including serotonergic 5-HT_1A_ in the raphe nuclei, dopaminergic D_2_ and DAT in the substantia nigra and ventral tegmental area, and noradrenergic NET in the locus coeruleus (Fig. S1). Next, we confirm that temporal signal-to-noise ratio (tSNR) in the brainstem, although low, is within the cortical tSNR range (Fig. S2a). Finally, we confirm that within the brainstem, smaller brainstem nuclei are not associated with lower tSNR (*r* = *−* 0.45, *p* = 0.0004, Fig. S2b, c).

**Figure 1.**
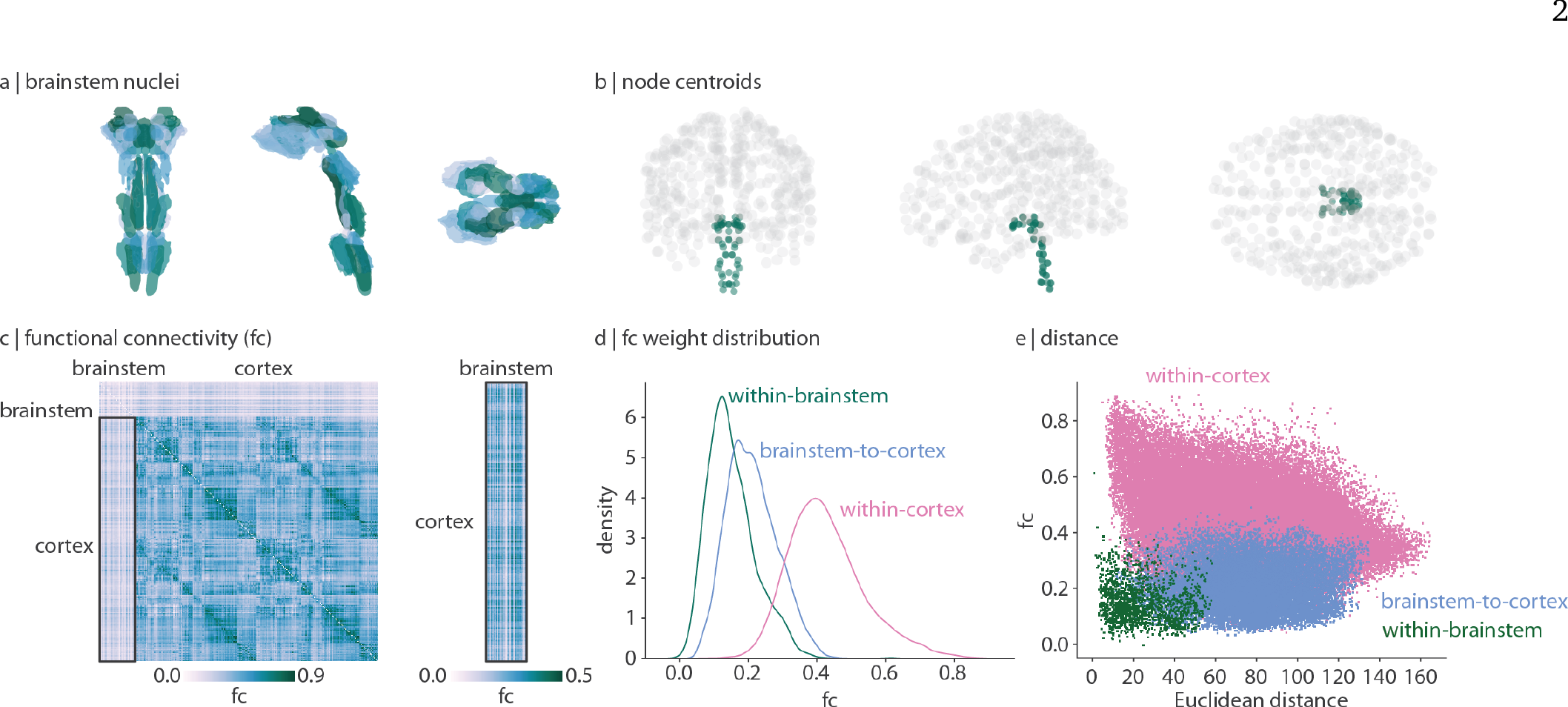
Brainstem-cortex functional connectivity. | (a) Coronal (posterior view), saggital, and axial view of the thresholded (35%) probabilistic template for all 58 brainstem nuclei in the Brainstem Navigator atlas (https://www.nitrc.org/projects/ brainstemnavig/ [16]). (b) Coronal (posterior view), saggital, and axial view of cortical (grey points, *n* = 400) and brainstem (green points, *n* = 58) parcel coordinate centroids. (c) Left: functional connectivity (FC) matrix (458 regions × 458 regions). Right: functional connectivity matrix between cortex and brainstem (400 cortical regions × 58 brainstem nuclei). (d) Density distributions of functional connectivity within brainstem (green), between brainstem and cortex (blue), and within cortex (pink). (e) Scatter plot of functional connectivity between regions as a function of Euclidean distance between parcel centroids. Within-cortex *r* = −0.29, *p* ≈ 0; brainstem-cortex *r* = 0.05, *p* = 8.7 × 10^*−*16^; within-brainstem *r* = −0.11, *p* = 3.4 × 10^*−*6^.

In Fig. 1c we show functional connectivity (FC; Pearson’s correlation between time-series) of the brainstem and cortex. Cortical FC shows a familiar network organization and is correlated with functional connectivity data from the Human Connectome Project (Spearman *r* = 0.58, *p* ≈ 0 [119]). Interestingly, we find that the brainstem is more functionally connected with the cortex than it is with itself (Fig. 1d, Welch’s twosided t-test *t* = 33.9, *p <* 0.001). Indeed, whereas cortical functional connectivity decreases with Euclidean distance [90], brainstem functional connectivity is less affected by distance (*r* = − 0.29 and *r* = − 0.11 respectively; Fig. 1e). This aligns with the intuition that brainstem nuclei primarily project to regions outside of the brainstem (including cortex, subcortex, and spinal cord), resulting in weak functional connectivity within the brainstem and stronger functional connectivity between brainstem and cortex.

### Dominant patterns of cortex-brainstem functional connectivity

The horizontal and vertical stripe patterning of brainstem-cortex FC shown in Fig. 1 indicates that there is a dominant pattern of brainstem-cortex connectivity. Hereafter we refer to the pattern of connectivity that brainstem nuclei make with the cortex as “brainstemto-cortex” connectivity, and vice versa for “cortex-to-brainstem” connectivity, despite no implication of directionality. The dominant pattern of how brainstem nuclei are connected with the cortex is quantified as the sum of FC across cortical regions (“weighted degree”, Fig. 2a). Brainstem-to-cortex hubs—brainstem nuclei that are most functionally connected with the cortex— are spatially segregated, in line with the theory that hub placement optimizes the trade-off between distance and efficient information transfer [26].

**Figure 2.**
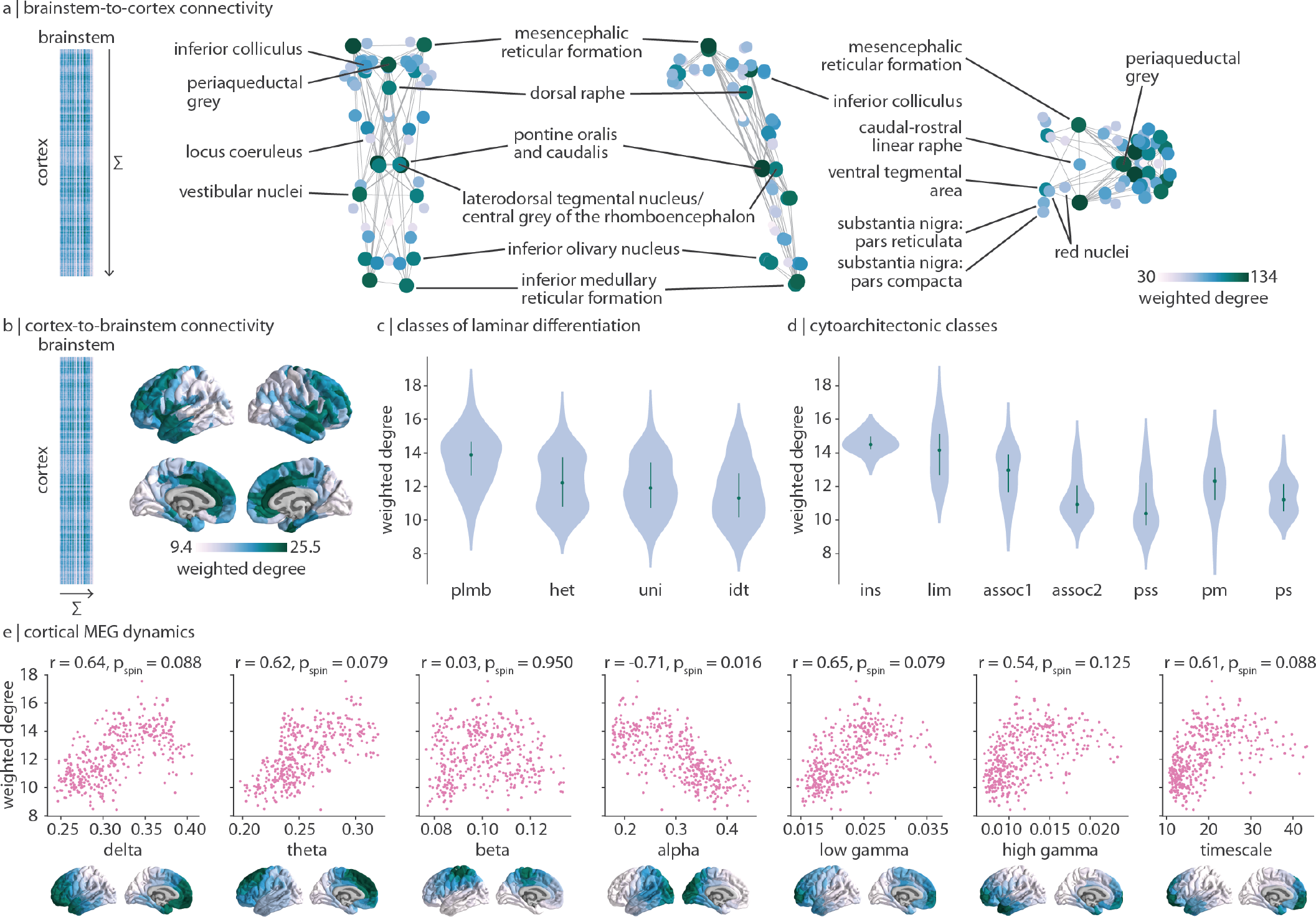
Dominant patterns of brainstem-cortex functional connectivity. | (a) Brainstem-to-cortex weighted degree is calculated by summing a brainstem nucleus’ functional connectivity across all cortical regions. Coronal (posterior view), sagittal, and axial perspectives of brainstem nuclei are shown. Node size and colour reflect weighted degree, and edges are plotted for the 5% strongest functional connections within the brainstem. Key brainstem nuclei are labelled. (b) Cortex-to-brainstem weighted degree is calculated by summing a cortical region’s functional connectivity across all brainstem nuclei. (c) Cortex-to-brainstem weighted degree binned according to classes of laminar differentiation (one-way ANOVA *F* = 18.5, *p* = 2.8 × 10^*−*11^) [69, 79]; plmb = paralimbic; het = heteromodal; uni = unimodal; idt = idiotypic. (d) Cortex-to-brainstem weighted degree binned according to classes of cytoarchitecture (one-way ANOVA *F* = 35.6, *p* = 2.0 × 10^*−*34^) [121, 122]; ins = insular; lim = limbic; assoc1 = association network 1; assoc2 = association network 2; pss = primary/secondary sensory; pm = primary motor; ps = primary sensory. (e) Scatter plots are shown for the correlation between cortex-to-brainstem weighted degree and seven metrics of MEG dynamics: power spectrum distributions for six canonical frequency bands and the intrinsic timescale (temporal memory of a neural element, see *Methods* for details); each point is a brain region (*N* = 400). Cortical distributions of MEG measures are shown on the brain surface below each plot and are derived from data in the HCP [119].

Brainstem-to-cortex hubs in the midbrain include the mesencephalic reticular formation, periaqueductal grey, and dorsal raphe [27, 103]. Brainstem hubs in the pons include the pontine reticular nuclei, the laterodorsal tegmental nucleus, and vestibular nuclei (spanning both pons and medulla). Finally, remaining brainstem hubs in the medulla include the inferior olivary nucleus and inferior medullary reticular formation. We confirm that the weighted degree pattern is not correlated with tSNR (Spearman *r* = 0.20, *p* = 0.14). We similarly show the weighted degree pattern in the cortex, which represents how strongly cortical regions are connected with the brainstem (cortex-to-brainstem hubs; Fig. 2b). This pattern follows an anterior-posterior gradient, with the anterior cingulate cortex being a primary hub of cortex-to-brainstem FC. This gradient also recapitulates classes of laminar differentiation (one-way ANOVA *F* = 18.5, *p* = 2.8 × 10^−11^ [69, 79]) and cytoarchitecture (one-way ANOVA *F* = 35.6, *p* = 2.0 × 10^−34^ [121, 122]), whereby limbic and insular regions demonstrate greatest brainstem functional connectivity while unimodal regions demonstrate the least brainstem connectivity (Fig. 2c, d).

The anterior-posterior cortical gradient of brainstem functional connectivity with cortex can be interpreted as a gradient of brainstem influence on cortical neural populations. Therefore, we tested whether this gradient is aligned with more direct measurements of cortical dynamics, that is, neural oscillatory rhythms from electrophysiology. Specifically, we correlate MEG-derived spectral power distributions for six canonical frequency bands as well as the intrinsic timescale (which can be interpreted as the temporal memory of a neural element) from the Human Connectome Project with cortexto-brainstem weighted degree [65, 98, 119]. We find that cortex-to-brainstem weighted degree is correlated (*r >* 0.5) with all seven measures of neural oscillatory dynamics, especially alpha power (which survives multiple comparisons correction and a spatial autocorrelation-preserving null; *r* = − 0.71, *p*_spin_ = 0.016; Fig. 2e). This demonstrates that cortical dynamics and brainstem input are aligned across multiple temporal resolutions.

### Brainstem connectivity reflects cognitive ontologies

Regions in the cortex and nuclei in the brainstem are all functionally connected with the whole brainstem following the same dominant pattern: the brainstem weighted degree pattern (originally shown in Fig. 2a; median *r* = 0.97, Fig. 3a left). To understand how brainstem nuclei are uniquely functionally connected with the cortex, we need to focus on connectivity patterns beyond this dominant pattern. We therefore regress brainstem weighted degree from each region’s connectivity-with-brainstem profile (Fig. 3a). This results in a functional connectivity matrix that represents how the brainstem and cortex are connected with one another above and beyond their dominant pattern of connectivity (Fig. 3a middle). By correlating the regressed cortical connectivity profile of pairs of brainstem nuclei, we construct a brainstem region × region correlation matrix that represents how similarly any two brainstem nuclei are functionally connected with the cortex (Fig. 3a right).

**Figure 3.**
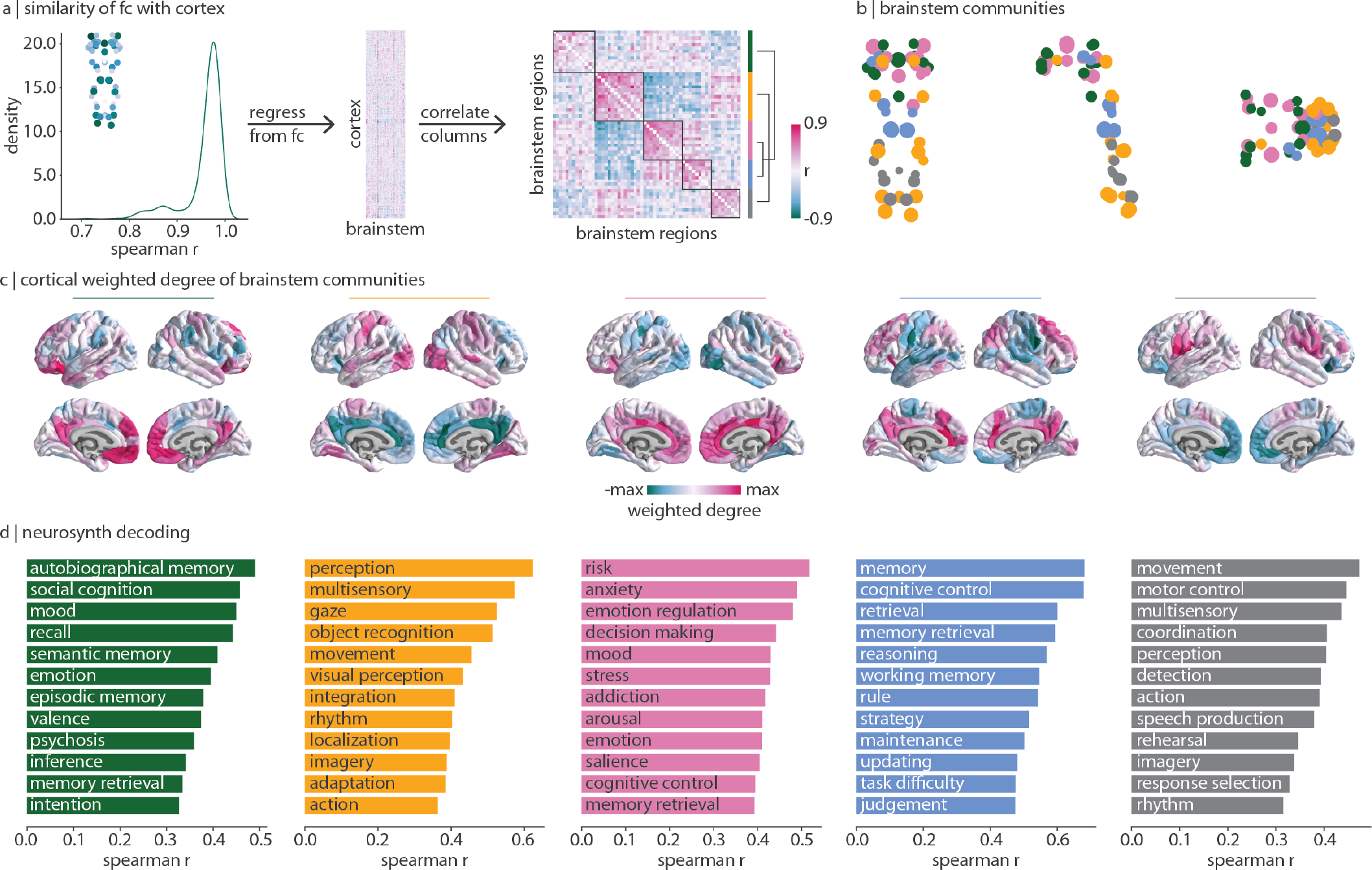
Brainstem communities underlying cortical function. | The Louvain community detection algorithm was applied to determine whether brainstem nuclei can be organized into distinct communities that make specific connectivity patterns with the cortex. (a) Left: for all 458 nodes (400 cortical, 58 brainstem), we correlate (Spearman *r*) the node’s brainstem functional connectivity profile with the weighted degree pattern shown in the inset and in Fig. 2a. The density distribution of Spearman’s *r* is shown (median *r* = 0.97). Middle: this brainstem map (weighted degree of brainstem-to-cortex functional connectivity) is regressed out of each cortical region’s brainstem functional connectivity pattern, resulting in a matrix (400 cortical regions × 58 brainstem nuclei) of functional connectivity residuals. Right: correlation matrix representing how similarly (Spearman’s *r*) two brainstem nuclei are functionally connected with the cortex, above and beyond the dominant pattern of connectivity between brainstem and cortex. Brainstem nuclei are ordered according to community affiliation (community colours shown on the right) and communities are outlined within the heatmap. Brackets on the right indicate how communities are joined in coarser community detection solutions (yellow combined with grey, blue combined with pink). (b) Community assignments from the Louvain community detection algorithm. Coronal (posterior view), sagittal, and axial perspectives of brainstem nuclei are shown. Node size is proportional to weighted degree shown in Fig. 2a. See Table 1 for a list of all brainstem nuclei organized by community affiliation. (c) Cortical weighted degree patterns are calculated as the sum of a cortical region’s functional connectivity with all brainstem nuclei within a specific community, and are shown for all five communities. These maps represent how each brainstem community is connected with the cortex. (d) Each cortical weighted degree pattern in panel (c) was correlated to 123 cognitive and behavioural meta-analytic activation maps from Neurosynth [126]. Only the top 10% correlations are shown.

This similarity matrix was subjected to the Louvain community detection algorithm at multiple resolution parameters (0.1 ≤ *γ* ≤ 6.0). We find that the brainstem can be divided into a nested hierarchy of communities, with each community representing a group of nuclei that exhibit similar functional connectivity patterns with the cortex. We show a stable solution of five approximately equally sized communities at *γ* = 2.8 in the main text (Fig. 3b) as well as two coarser solutions in the supplement which emerge when combining communities (Fig. S3, S4). Regions within each community are listed in Table 1 and we describe each community in detail below. How are these brainstem communities connected with the cortex? For each brainstem community, we calculate each cortical region’s total functional connectivity (weighted degree; sum of FC across brainstem nuclei) with all brainstem nuclei within a specific community (Fig. 3c). This results in a cortical network pattern that is associated with each brainstem community. Next, to determine the functional specialization of each cortical network, we correlate the cortical weighted degree patterns in Fig. 3c with 123 meta-analytic functional activation patterns from Neurosynth (see *Methods* for details [126]). We show the 12 (10%) most highly correlated Neurosynth keywords in Fig. 3d.

**TABLE 1.** Brainstem communities. | Brainstem nuclei within each of the five communities shown in Fig. 3. Asterisks indicates a midline nucleus. L/R refers to the hemisphere of bilateral nuclei.

| green | yellow | pink | blue | grey |
| --- | --- | --- | --- | --- |
| median raphe nucleus* | raphe obscurus* | dorsal raphe* | isthmus reticular formation (L) | raphe magnus* |
| paramedian raphe nucleus* | raphe pallidus* | caudal-rostral linear raphe* | pontine reticular nucleus: pontis oralis and caudalis (LR) | parvicellular reticular nucleus: alpha part (L) |
| periaqueductal grey* | inferior olivary nucleus (LR) | ventral tegmental area/parabrachial pigmented nucleus complex (LR) | laterodorsal tegmental nucleus/central grey of the rhombencephalon (LR) | superior medullary reticular formation (LR) |
| substantia nigra: pars reticulata (L) | lateral parabrachial nucleus (LR) | median parabrachial nucleus (R) | microcellular tegmental nucleus-parabigeminal nucleus (L) | superior olivary complex (LR) |
| substantia nigra: pars compacta (LR) | inferior medullary reticular formation (LR) | substantia nigra: pars reticulata (R) | median parabrachial nucleus (L) | viscero-sensory-motor nuclei complex (LR) |
| red nucleus: subregion 1 (LR) | parvicellular reticular nucleus: alpha part (R) | mesencephalic reticular formation (LR) | locus coeruleus (LR) | subcoeruleus (L) |
| red nucleus: subregion 2 (R) | red nucleus: subregion 2 (L) | cuneiform (LR) |  |  |
| pedunculoegmental nuclei (L) | vestibular nucleus (LR) | isthmus reticular formation (R) |  |  |
| microcellular tegmental nucleus, parabigeminal nucleus (R) | subcoeruleus (R) | pedunculotegmental nucleus (R) |  |  |
| superior colliculus (LR) | inferior colliculus (LR) |  |  |  |

We find a community (yellow) composed of regions throughout the brainstem including the inferior colliculus, vestibular nuclei, and inferior olivary nucleus. This community is most functionally connected with unimodal cortex and associated with sensory perception and movement. A second sensory-related community (grey) exists in the medulla and is composed of regions including the superior olivary complex, the viscero-sensorymotor complex, and the raphe magnus. This community is most connected with ventral regions of primary motor and sensory cortex, as well as anterior parietal regions such as the angular and supramarginal gyri, regions that are associated with higher-order motor coordination and speech. Note that the yellow and grey communities are joined in the three-community solution (Fig. S3). We also find a community (pink) composed of midbrain regions including the ventral tegmental area, dorsal and caudal-rostral linear raphe nuclei, and mesencephalic reticular formation. This community is most functionally connected with cingulate and limbic regions associated with emotion regulation, affect, addiction, and arousal.

Finally, we find two brainstem communities that are related to higher-order cognitive functions. The first (green) is composed of midbrain regions including the substantia nigra, red nucleus, superior colliculus, and periaqueductal grey. This community is most connected with medial transmodal cortical regions including the precuneus and frontal pole. The second higher-order cognitive community (blue) is composed of regions in the midbrain and pons including the locus coeruleus, the laterodorsal tegmental nucleus/central grey of the rhomboencephalon, and the pontine reticular nuclei. Both the green and blue communities are functionally connected with transmodal cortex, and are associated with memory, but each community is specialized. The green community is most connected with the frontal pole and is associated with autobiographical memory and social cognition. Meanwhile, the blue community is connected more broadly to medial and lateral transmodal cortex and is associated with memory retrieval, working memory, and cognitive control. Notably, the green community remains isolated in the three- and four-community solutions, while the blue and pink communities are combined (Fig. S3, S4). Altogether, this finding demonstrates the striking alignment between cognitive function and brainstem function.

### Mapping chemoarchitecture to brainstem communities

Since the cortex receives input from multiple neuromodulatory brainstem nuclei, we sought to identify the relationship between neurotransmitter systems, the identified brainstem communities, and their cortical projection patterns. We used data from a recently acquired PET atlas of nine neurotransmitter systems in the human brain to estimate cortical distributions of 18 neurotransmitter receptors and transporters [51, 65]. Specifically, for each brainstem community, we fit a multiple linear regression model that predicts the community’s cortical weighted degree profile from receptor and transporter densities (Fig. 4a). Next, we apply dominance analysis to estimate the relative contribution (“dominance”) of each receptor and transporter to the overall fit (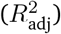) of the model (Fig. 4b) [4]. The norepinephrine transporter (NET) emerges as a dominant receptor across all communities, peaking in the blue memory community which includes the primary nucleus for norepinephrine synthesis: the locus coeruleus. The second higher-order cognitive brainstem community (green) is connected with the cortex in the manner that aligns with monoamine transporters including dopaminergic DAT and serotonergic 5-HTT. Indeed, this community includes the dopaminergic substantia nigra and serotonergic median and paramedian raphe nuclei.

**Figure 4.**
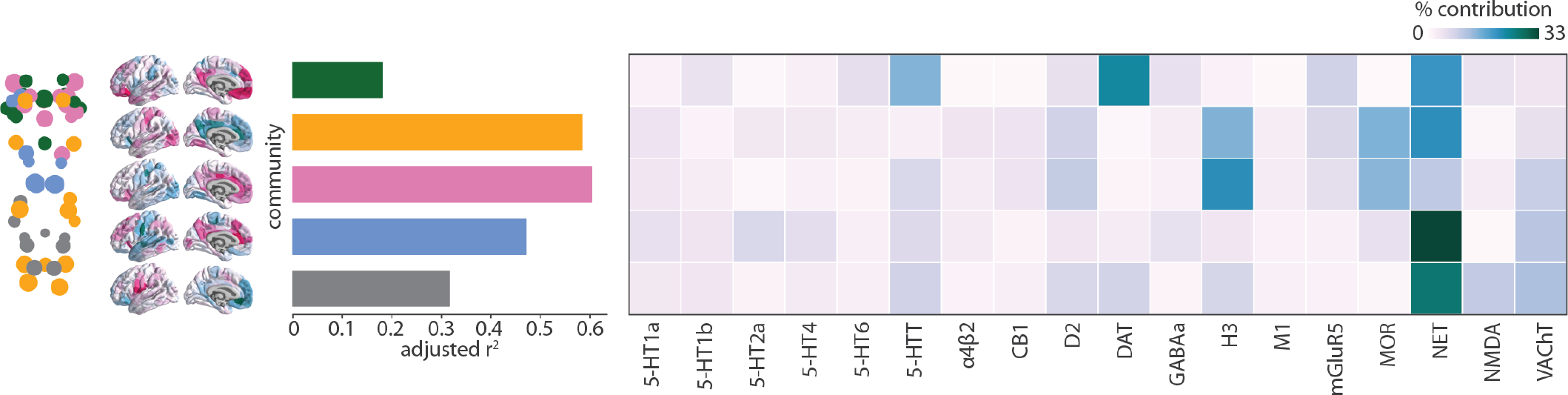
Mapping chemoarchitecture to brainstem communities. | For each community (shown on the brainstem plot on the left, as well as in Fig. 3b), a multiple linear regression model was fit between 18 cortical neurotransmitter receptor and transporter density profiles and the community’s cortical weighted degree pattern (shown as surface plots, as well as in Fig. 3c). Model fits (adjusted *R*^2^) are shown in the bar plot. Dominance analysis was applied to the independent variables (receptors and transporters) to determine which receptors/transporters were contributing most to the model fit [4] Percent contribution is shown in the heatmap. Receptor/transporter data were acquired from a PET atlas of neurotransmitter receptor/transporter densities in the human brain [51, 65].

We find that 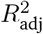 is greatest for the sensory (yellow) and affect (pink) brainstem communities. In other words, these brainstem nuclei are functionally connected with the cortex in a manner that is more aligned with cortical receptor distributions than other brainstem communities (Fig. 4a). The most dominant receptors for both these communities include the histamine receptor H_3_, opioid receptor MOR, norepinephrine transporter NET, dopamine receptor D_2_, and for the pink community also serotonin transporter 5-HTT and acetylcholine transporter VAChT. These receptors span multiple neurotransmitter systems and are primarily metabotropic rather than ionotropic. Collectively, our findings highlight the role that multiple transmitter systems play in modulating brainstem-cortex functional connectivity.

### Brainstem nuclei delineate unimodal and transmodal cortex

Lastly, we ask: if brainstem nuclei demonstrate unique patterns of functional connectivity with the cortex, do cortical regions likewise demonstrate unique patterns of functional connectivity with the brainstem? Using the regressed functional connectome described above, we correlate the regressed brainstem connectivity profile of pairs of cortical regions to construct a cortical region × region correlation matrix that represents how similarly two cortical regions are functionally connected with the brainstem (Fig. 5a). Interestingly, by exploring functional connectivity profiles between cortex and brainstem, we find that cortical regions are aligned with cortical intrinsic functional networks [127] (Fig. 5a right).

**Figure 5.**
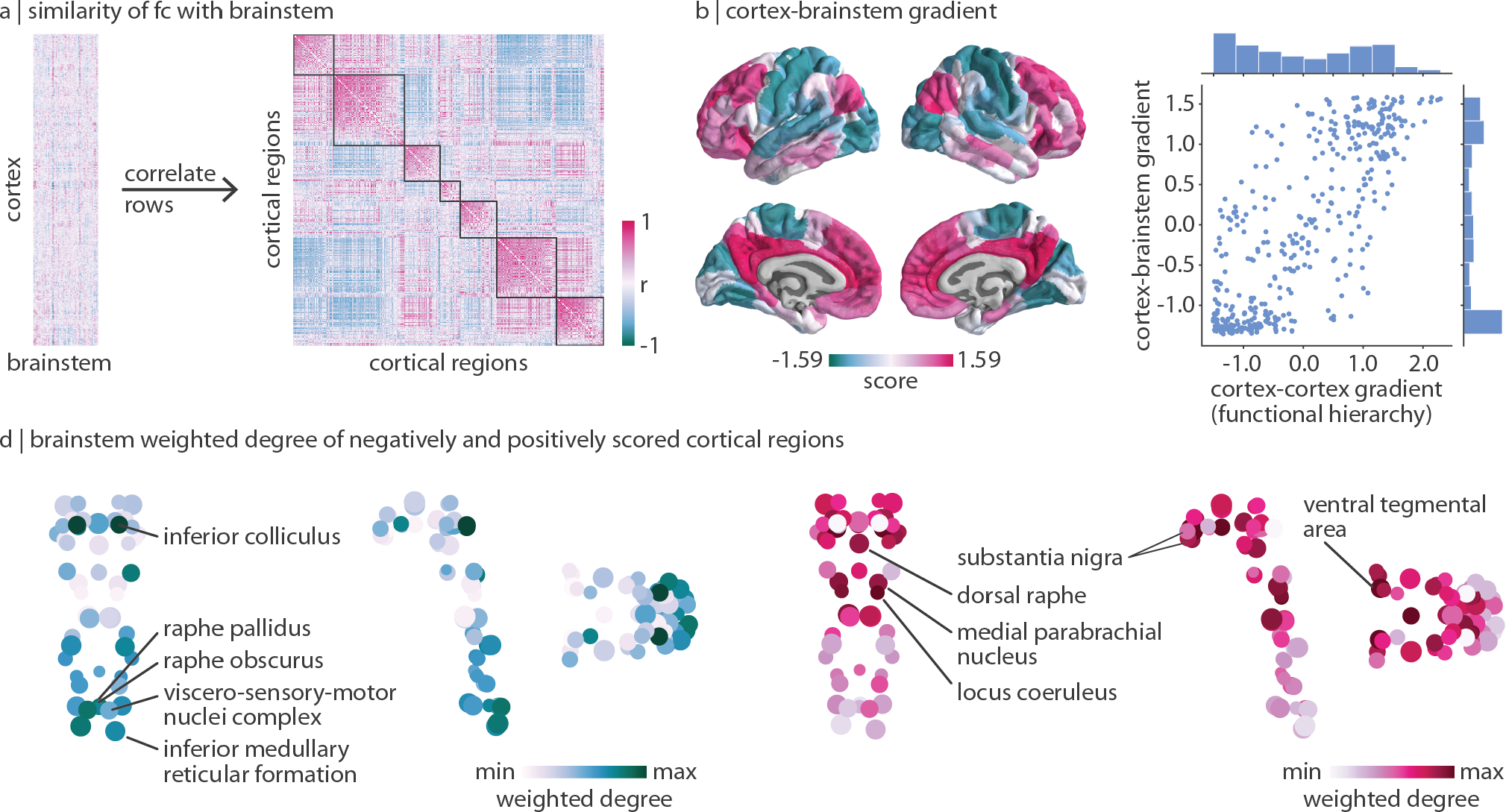
Brainstem nuclei delineate unimodal and transmodal cortical regions. | (a) Left: functional connectivity residuals (identical to the matrix shown on the left in Fig. 3a). Right: correlation matrix represents how similarly (Spearman’s *r*) two cortical regions are functionally connected with the brainstem above and beyond the dominant pattern of brainstem-cortex connectivity. Outlines are shown around the seven Yeo-Krienen resting-state networks (order: control, default mode, dorsal attention, limbic, ventral attention, somato-motor, visual). (b) Diffusion map embedding was applied to the matrix shown in panel (a). Left: the first gradient of cortex-brainstem functional connectivity. Right: correlation between the first gradient of cortex-brainstem connectivity and the first gradient of cortex-cortex functional connectivity (also called the cortical functional hierarchy, the unimodal-transmodal axis, and the sensory-association axis; *r* = 0.77, *p*_spin_ = 0.0001). Distribution of gradient values are shown for both gradients. (d) Brainstem weighted degree patterns are calculated as the sum of a brainstem nucleus’ functional connectivity with all negatively- (left) or positively- (right) scored regions of the cortical gradient shown in panel (b). Coronal (posterior view), sagittal, and axial perspectives of brainstem nuclei are shown. Node size is proportional to weighted degree shown in Fig. 2a.

Furthermore, we use diffusion map embedding to calculate the first gradient of how similarly cortical regions are connected with the brainstem. Cortical regions with similar scores along this gradient are similarly connected with the brainstem; the greater the difference in gradient scores, the more dissimilar regions are in their brainstem connectivity profiles. This gradient is strongly correlated with the principal functional gradient of corticocortical connectivity (also derived using diffusion map embedding; *r* = 0.77, *p*_spin_ = 0.0001; Fig. 5b), which is thought to delineate a hierarchy of cortical function from unimodal (e.g. primary regions involved in lower-order functions) to transmodal (e.g. association regions involved in higher-order functions) regions [64, 69]. We find that this cortical unimodal-transmodal hierarchy also reflects brainstem functional connectivity. Interestingly, for the gradient derived from cortical connectivity with the brainstem, most cortical regions are placed at the extremes of the gradient. Indeed, the most stable solution from the Louvain community detection algorithm is one that identifies two prominent communities (one transmodal, one unimodal) (Fig. S5).

Which brainstem nuclei are more functionally connected with unimodal (negative gradient score) and transmodal (positive gradient score) regions? We calculate the weighted degree of FC from negatively- and positively-scored cortical regions to the brainstem (Fig. 5d). We find that unimodal brain regions are most connected with caudal brainstem nuclei in the medulla including the inferior medullary reticular formation, the viscero-sensory-motor nuclei complex, and the raphe pallidus and obscurus. In addition to these nuclei, the brainstem nucleus with the greatest unimodal connectivity is the inferior colliculus in the midbrain. Likewise, brainstem nuclei most connected with transmodal regions exist in the midbrain and pons including the ventral tegmental area, the locus coeruleus, the substantia nigra, the dorsal raphe, and the medial parabrachial nucleus [8]. Altogether, we find that specific brainstem nuclei are connected with unimodal and transmodal cortex, an indication that cortical hierarchies are not derived only from cortical function.

### Sensitivity, robustness, and replication in subcortex

We conduct three analyses to gauge the sensitivity and robustness of the current findings. First, we run a splithalf resampling analysis where we randomly split the sample of 20 individuals into two groups of 10 (100 repetitions). We correlate the group-averaged functional connectomes, brainstem-to-cortex hub profile, and cortex-to-brainstem hub profile between the two groups to assess how much results vary given different samples of participants. We find functional connectivity (0.85 *< r <* 0.95), brainstem-to-cortex hubs (0.90 *< r <* 1), and cortex-to-brainstem hubs (0.7 *< r <* 0.92) are all highly correlated between groups (Fig. S6). Next, to ensure results are robust against alternative parcellations, we repeat the analyses using a 100-region cortical parcellation [96] and find consistent results for all analyses (Fig. S7). Third, since all participants underwent both 7 Tesla and 3 Tesla scanning, we were able to reconstruct their functional connectomes from the 3 Tesla fMRI time-series. When comparing the functional connectomes across these two scanning conditions, we find that within-cortex functional connectivity is correlated at *r* = 0.40, within-brainstem functional connectivity is correlated at *r* = 0.75, and brainstem-to-cortex functional connectivity is correlated at *r* = 0.70 (Fig. S8). Altogether, these analyses demonstrates that our findings are generalizable across different scanners and processing pipelines.

Finally, for completeness, we extend the analyses to other subcortical and diencephalic regions—physically located between the cortex and brainstem and likely mediating their relationship—as a first step towards understanding how the present findings are reflected in subcortical structures (Fig. S9a). Specifically, 7 Tesla functional data was also acquired for 14 bilateral FreeSurfer subcortical regions (caudate, putamen, pallidum, nucleus accumbens, thalamus, amygdala, hippocampus [38]), 8 bilateral Brainstem Navigator diencephalic nuclei (lateral geniculate nucleus (LGN), medial geniculate nucleus (MGN), subthalamic nuclei subregions 1 & 2 [16]), and the hypothalamus [80]. We refer to the FreeSurfer structures as “subcortex” (although the hippocampus is technically allocortex, and the thalamus is also part of the diencephalon) and the Brainstem Navigator diencephalic nuclei plus hypothalamus as “diencephalon”. Importantly, the FreeSurfer subcortical regions are large, cytoarchitectonically defined brain regions and do not undergo brainstem-specific preprocessing, whereas the diencephalic nuclei are small nuclei defined from T1 images and do undergo brainstem-specific processing due to their size and proximity to vasculature and CSF.

Fig. S9b shows the weighted degree of brainstem functional connectivity of subcortical and diencephalic nuclei. Regions with the greatest functional connectivity with the brainstem include the thalamus, hypothalamus, and the LGN, supporting the notion that the thalamus is a main hub of cortex-subcortex connectivity [102]. In Fig. S9c, we reconstruct the region × region correlation matrix representing how similarly non-neocortical regions are functionally connected with the neocortex, above and beyond the dominant weighted degree pattern of connectivity. We apply the Louvain community detection and find a similar modular decomposition of the brainstem as in the main analyses. Finally, we apply diffusion map embedding to the region × region similarity matrix representing how similarly neocortical regions are functionally connected with non-neocortical structures. We find that the first gradient still resembles the unimodal-transmodal gradient (Fig. S9d), and we find that the hippocampus, amygdala, and MGN have greatest functional connectivity with negatively-scored (unimodal) brain regions while the caudate, putamen, thalamus, and LGN have greatest functional connectivity with positively-scored (transmodal) brain regions. Altogether, the present findings remain consistent when extended to subcortical regions and diencephalic nuclei. An in-depth study of the relationship between cortical, subcortical, and brainstem functional architectures is left for future work.

## DISCUSSION

In the present report, we use a high resolution 7 Tesla fMRI dataset in conjunction with a comprehensive brainstem atlas of 58 nuclei to investigate how cortical function reflects brainstem function. We identify a compact set of integrative hubs in the brainstem with strong functional connectivity with the cortex. We find that multiple cortical phenomena, including oscillatory rhythms, cognitive function, and the unimodal-transmodal hierarchy, can be traced back to specific functional connectivity with the brainstem.

*In vivo* functional imaging of the human brainstem has long eluded the neuroimaging field due to the challenges of imaging this constellation of deep structures, resulting in a vacuum of knowledge about awake human brainstem activity [97]. In the last decade, substantial progress has been made to acquire a robust functional signal in the brainstem. Extensive research on the sources of fMRI signal in the brainstem and the necessity of noise correction has improved the acquisition and interpretability of signals in deep brain structures [14, 18, 23]. In 2015, Bianciardi et al. [16] began developing an *in vivo* neuroimaging template of human brainstem nuclei which facilitated the standardization of whole-brainstem functional imaging. Later in 2022, Singh et al. [103] and Cauzzo et al. [27] reported the resting state functional connectomes of arousal and motor [103] and autonomic, limbic, pain, and sensory [27] brainstem nuclei to the rest of the brain. In the present study, we join these connectomes into a single dataset of whole-brainstem to whole-cortex functional connectivity to ask: what can *in vivo* human whole-brainstem functional activity tell us about cortical function?

First we locate the regions in the brainstem that are most functionally connected with the cortex. While there exists a rich literature of hubs in cortex [25], little is known about the hubs in the brainstem [8, 27, 103]. We identify a set of integrative brainstem hubs that are located throughout the midbrain, pons, and medulla. Brainstem-to-cortex hubs are functionally diverse, with some thought to be primarily involved in motor functions (e.g. inferior olivary nucleus (motor coordination), pontine nuclei (movement), vestibular nuclei (balance)), some that are associated with specific neurotransmitter systems (e.g. dorsal raphe (serotonin), laterodorsal tegmental nucleus (acetylcholine) [118]), and some that have been linked to multiple functions (e.g. mesencephalic and inferior medullar reticular formation, periaqueductal grey). Surprisingly, the locus coeruleus is not identified as a hub, despite its known widespread projections throughout the cortex and its role in information integration [123]. However, previous work has speculated that the locus coeruleus’ integrative properties only emerge during specific behavioural contexts [22]. This suggests that brainstem hubs may be state-dependent and temporally variable—an exciting direction for future work.

Likewise, we demonstrate that cortical regions follow an anterior-posterior gradient with respect to their functional connectivity strength to the brainstem, with the largest cortex-to-brainstem hubs existing in anterior cingulate cortex. Previous studies have reported greater diffusion-weighted MRI-derived structural connectivity from anterior cortex to brainstem [44, 105]. Similarly, transcriptomic analysis of human von Economo neurons, whose cell bodies are restricted to layer V of the anterior cortex, have shown that these bipolar neurons express transcriptional factors associated with long-range projections to the brainstem [30, 57]. In other words, this gradient of cortex-brainstem functional connectivity likely reflects the underlying synaptic connectivity between brainstem and cortex [111]. Finally, we find a close correspondence between cortex-to-brainstem hubs and MEG-derived alpha power. Although cortical rhythms have been extensively studied, subcortical and brainstem rhythms are difficult to measure because of electrophysiological signal decay across larger distances [5]. Our work suggests that brainstem connectivity informs the grammar of ongoing cortical dynamics, prompting future work to test the relationship between cortical and brainstem rhythms [68, 114]. Altogether, the anteriorposterior cortical gradient of brainstem connectivity reflects both brain structure and oscillatory dynamics.

Functional imaging has been used to demonstrate networks of cortical regions that coactivate both during specific tasks and at rest [40, 109, 127]. Recent studies that explore extracortical structures of the central nervous system have demonstrated that cortical networks also coactivate with specific cerebellar regions [24, 50, 110], spinal cord segments [117], and specific brainstem nuclei [8, 11]. Rather than imposing cortically-defined patterns of functional activation on the brainstem, we asked whether brainstem nuclei share similar connection patterns with the cortex; and if so, what are the cortical networks of brainstem connectivity? We find that the brainstem can be organized into hierarchical communities of nuclei with similar cortical connectivity. These communities establish functional links between brainstem nuclei that were previously unknown, and can likely only be observed in living humans where it is possible to record neural activity simultaneously from the brainstem and cortex. Furthermore, each brainstem community is connected with familiar cortical functional networks underlying cognition, memory, sensation, movement, and emotion. This indicates that the brainstem is not simply a structure for executing evolutionarily conserved functions, but rather that the brainstem has widespread involvement in multiple cognitive and behavioural functions [8, 60].

Each community is associated with specific patterns of functional specialization. A natural explanation is that this is due to their specific chemoarchitectural makeup. Namely, the brainstem is made up of multiple neuromodulatory systems which project throughout the cortex, tuning large-scale synchronization of neuronal populations and emergent functions [101, 118]. A major neuromodulatory system that projects throughout the brain is the noradrenergic system [85, 115]. We find that the norepinephrine transporter is closely aligned with each brainstem community’s associated cortical activation pattern, and that this relationship is strongest in the community (blue) which houses the noradrenergic locus coeruleus (Fig. 3). Furthermore, this community is related to memory, cognitive control, and retrieval, all integrative functions thought to be controlled by the norepinephrine system [100]. We also find a community (green) composed of the dopaminergic substantia nigra and the serotonergic median raphe nucleus, and underlying memory and social cognition, that is most aligned with dopaminergic and serotonergic transporters. Ultimately, by integrating neurotransmitter receptor/transporter datasets with simultaneous brainstem-cortex fMRI, we find evidence that neuromodulatory systems help shape cortical-brainstem interactions.

Finally, we find that cortical regions are connected with the brainstem following a well known and frequently studied cortical gradient: the sensory-association axis [64, 69, 112]. The sensory-association axis, or unimodal-transmodal functional hierarchy, describes a gradient of cortical function from lower- to higher-order processes. This gradient is aligned with cortical expansion across ontogeny [55] and phylogeny [55, 125], becomes more polarized with development [63, 83], and less polarized with pathological progression [54]. Notably, the sensory-association axis is generally observed from and interpreted in light of cortical processing and cortico-cortical connectivity. Here we find that the poles of the sensory-association axis demonstrate distinct connectivity patterns with the brainstem. This may indicate that functional inputs from the brainstem anchor the polar extremes of the cortical hierarchy (i.e. primary and association cortex), while cortico-cortical connectivity patterns fill in the gradual shift from lowerto higherorder cortical functions. In other words, the hierarchy of cortical function may emerge from connectivity patterns with the brainstem, bringing to light the influence that extracortical structures can have on cortico-cortical connectivity. How the brainstem is involved in gradient changes across development, healthy aging, and pathology is an exciting question for future research.

In the present report, we extend our umwelt of *in vivo* cortical functional networks to the brainstem and find that multiple cortical phenomena are reflected by brainstem-cortex functional connectivity. This opens doors for many future applications of brainstem functional connectivity. For example, multiple pathological markers, such as *α*-synuclein in Parkinson’s disease, are thought to emerge from brainstem dysfunction before spreading throughout the cortex [21, 58]. Brainstem functional connectivity patterns may generate more accurate models of disease propagation and aberrant dynamics, giving rise to potentially actionable brainstem targets [129]. Brainstem functional connectivity may also facilitate the development of better computational models of ongoing dynamics [70, 124]. While the present work extends the study of *in vivo* human cortical function to the brainstem, it is increasingly possible to integrate not only brainstem function, but the structure and function of the cerebellum, subcortex, and spinal cord, into a single wiring diagram of the complete human central nervous system [44, 49, 53, 105].

We close with some important methodological considerations. First, brainstem nuclei are notoriously difficult to image, given their deep location, proximity to vasculature and CSF, their irregular shape, and small size. This brainstem dataset underwent extensive and optimized physiological noise correction and validation of the defined nuclei, but brainstem imaging is an active area of research and best practices continue to be refined. Second, the temporal resolution of the 7 Tesla fMRI was minimized at 2.5 seconds. This was necessary given the number of slices and spatial resolution required to reconstruct small brainstem nuclei. Third, only 20 healthy participants were included in this study. Although we replicate the findings using 3 Tesla scans in the same participants and perform a split-half resampling analysis, future work is necessary to validate our findings in largeN datasets. Fourth, the optimal brainstem registration may result in suboptimal cortical registration, although we find that within-cortex FC is correlated with FC from an independent dataset (Human Connectome Project).

In summary, we map the functional architecture of brainstem-cortex connectivity. We find that the functional architecture of the brainstem is an ever-present leitmotif of cortical function. The present work takes advantage of advances in modern brain imaging, extending the scope of inquiry to structures that were previously inaccessible, and ultimately leading to a more complete understanding of the brain.

## METHODS

All preprocessed data and code used to perform the analyses are available at https://github.com/netneurolab/hansen_brainstemfc.

### fMRI data acquisition

Functional magnetic resonance imaging (fMRI) data in the brainstem was collected, preprocessed, and originally presented by Cauzzo et al. [27] and Singh et al. [103]. 20 unrelated healthy participants (age range 29.5 *±* 1.1 years, 10 males, 10 females) participated in two eyes-closed resting-state 7 Tesla and 3 Tesla MRI sessions (Magnetom and Connectom respectively, Siemens Healthineers, Erlangen, Germany). During the 7 Tesla session, three runs of 10 minutes were acquired, while a single run of 9 minutes was acquired at 3 Tesla. Notably, brainstem-specific custom protocols were developed for the 7 Tesla MRI acquisition and processing, which we describe below, whereas conventional sequences were used for the 3 Tesla MRI acquisition. Complete details for all acquisition and processing parameters are detailed in full in both Cauzzo et al. [27] and Singh et al. [103].

Briefly, a custom-built 32-channel receive coil and volume transmit coil was used at 7 Tesla, and a custom-built 64-channel receive coil and volume transmit coil was used at 3 Tesla. For each subject, three runs of 7 Tesla functional gradient-echo echo-planar images (EPIs) were acquired with the following parameters: isotropic voxel size = 1.1 mm, matrix size = 180 *×* 240, GRAPPA factor = 3, nominal echo-spacing 0.82 ms, bandwidth = 1488 Hz/Px, number of slices = 123, slice orientation = sagittal, slice-acquisition order = interleaved, echo time (TE) = 32 ms, repetition time (TR) = 2.5 s, flip angle (FA) = 75^°^, simultaneous-multi-slice factor = 3, number of repetitions = 210, phase-encoding direction = anterior-posterior, acquisition-time = 10 minutes, 7 seconds. Between the three fMRI runs, subjects’ awake state was verified verbally. Foam pads were used to minimize head motion and earplugs were provided. To account for physiology related signal fluctuations, timing of cardiac and respiratory cycles was recorded via piezoelectric finger pulse sensor (ADInstruments, Colorado Springs, CO, USA) and piezoelectric respiratory bellow (UFI, Morro Bay, CA, USA), respectively. To correct for geometric distortion, a 2.0 mm isotropic resolution fieldmap was acquired.

### fMRI data preprocessing

Physiological noise correction was done in each resting state fMRI run using custom-built Matlab function of RETROICOR [48] adapted to the slice acquisition sequence. Functional images were then slice-time corrected, reoriented to standard orientation, and coregistered to the MEMPRAGE image. Coregistration was implemented in AFNI using a two-step procedure made of an affine coregistration and a boundary-based (edge enhancing) nonlinear coregistration [31]. Next, six rigid-body motion time-series nuisance regressors, a regressor describing respiratory volume per unit time convolved with a respiration response function [19], a regressor describing heart rate convolved with a cardiac response function [28], and five regressors modeling the signal in cerebrospinal fluid (CSF), extracted using PCA on a mask of the ventricles, were regressed from the fMRI time-series. Cleaned data were scaled to percent signal change by dividing by the temporal signal mean, multiplying by 100, and bandpass filtering between 0.01–0.1 Hz. Finally, any residual temporal mean was removed and the three runs were concatenated.

### Functional network reconstruction

To construct a 2D functional connectome for each participant, we used the 400-region Schaefer atlas in the cortex [96] and the Brainstem Navigator atlas in the brainstem (https://www.nitrc.org/projects/brainstemnavig/ to define seed and target regions [15, 16, 45, 46, 104, 106]). The Brainstem Navigator is a probabilistic atlas of 58 brainstem nuclei throughout the midbrain, pons, and medulla (50 bilateral nuclei, 8 midline nuclei), aligned to the MNI template. In all analyses we threshold the probabilistic atlas at 35%. Since brainstem nuclei vary in size (quantified as the number of voxels within each region), we confirm that parcel size does not reflect temporal signal-to-noise ratio (tSNR) (Fig. S2). tSNR was calculated as the mean of the time-series divided by the standard deviation (before demeaning the time-series in the preprocessing steps outlined above), averaged across subjects, and parcellated to the defined cortical and brainstem regions. Finally, functional connectivity was defined as the Pearson’s correlation between time-series for every pair of brain regions (458 total). The group-averaged connectome was calculated as the mean across individual subject connectomes. Analyses were repeated using a 100-region Schaefer atlas as part of the robustness analysis. The final functional connectivity matrix was compared with a standard 3T functional connectivity matrix from the HCP (326 unrelated participants, age range 22—35 years, 145 males, S900 release), downloaded from https:// github.com/netneurolab/hansen_many_networks [52].

### MEG data acquisition and preprocessing

6-minute resting state eyes-open magenetoencephalography (MEG) time-series were acquired from the Human Connectome Project (HCP, S1200 release) for 33 unrelated participants (age range 22—35, 17 males) [47, 119]. Complete MEG acquisition protocols can be found in the HCP S1200 Release Manual. For each participant, we computed the power spectrum at the vertex level across six different frequency bands: delta (2–4 Hz), theta (5–7 Hz), alpha (8–12 Hz), beta (15–29 Hz), low gamma (30–59 Hz), and high gamma (60–90 Hz), using the open-source software, Brainstorm [113]. The preprocessing was performed by applying notch filters at 60, 120, 180, 240, and 300 Hz, and was followed by a high-pass filter at 0.3 Hz to remove slow-wave and DC-offset artifacts. Preprocessed sensor-level data was used to obtain a source estimation on HCP’s fsLR4k cortex surface for each participant. Head models were computed using overlapping spheres and the data and noise covariance matrices were estimated from the resting state MEG and noise recordings. Brainstorm’s linearly constrained minimum variance (LCMV) beamformers method was applied to obtain the source activity for each participant. Welch’s method was then applied to estimate power spectrum density (PSD) for the source-level data, using overlapping windows of length 4 seconds with 50% overlap. Average power at each frequency band was then calculated for each vertex (i.e. source). Source-level power data was then parcellated into 400 and 100 cortical regions for each frequency band, according to the Schaefer atlas [96]. Intrinsic timescale of the MEG signal was estimated using spectral parameterization with the FOOOF (fitting oscillations & one over *f*) toolbox [33], via the method developed by Gao et al. [43]. The intrinsic timescale map for the HCP dataset was first calculated and analyzed in Shafiei et al. [99]. All pre-processed brain maps were downloaded directly from neuromaps [65].

### Community detection

To identify communities of brainstem nodes that are similarly connected with the cortex, we applied the Louvain community detection algorithm [20]. Since both brainstem and cortex are connected with the brainstem following a dominant pattern (Fig. 2a), we first regressed this weighted degree pattern from every node’s connectivity-to-brainstem profile (Fig. 3a). The residuals represent the degree to which nodes are connected with one another above and beyond this dominant pattern of connectivity. Second, we constructed a brainstem region × brainstem region similarity matrix by correlating the cortical connectivity profiles of every pair of brainstem nodes. This similarity matrix was subjected to the Louvain algorithm, which maximizes positive correlations within communities and negative correlations between communities.

Specifically, brainstem nodes were assigned to communities in a manner that maximizes the quality function

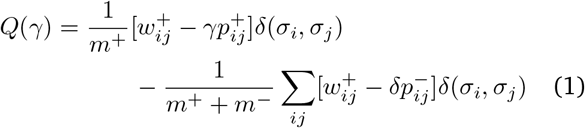

where 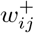 is the network with only positive correlations and likewise for 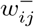 and negative correlations. The term 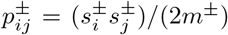 represents the null model: the expected density of connections between nodes *i* and *j*, where 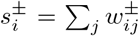 and 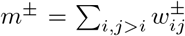. The variable *σ*_*i*_ is the community assignment of node *i* and *δ*(*σ*_*i*_, *σ*_*j*_) is the Kronecker function and is equal to 1 when *σ*_*i*_ = *σ*_*j*_ and 0 otherwise. The resolution parameter, *γ*, scales the relative importance of the null model, wither greater *γ* (*γ >* 1) making it more difficult to detect large communities. In other words, as *γ* increases, increasingly fine network partitions, and more communities, are identified. We tested 60 values of *γ*, from *γ* = 0 .1 to *γ* = 6.0, in increments of 0.1. At each *γ*, we repeated the algorithm 250 times and constructed a consensus partition, following the procedure recommended in Bassett et al. [9].

For each *γ*, the similarity of the clustering solution across the 250 partitions was calculated as the z-score of the rand index. Consensus partitions are considered better quality (i.e. more stable) when the mean of the z-scored rand index is high and the variance is low. We show the mean and variance of the z-scored rand index across all *γ*, as well as the number of communities identified, i n Fig. S 10. We show the community solution at *γ* = 2.8 in the main text because it identifies approximately equally sized communities (Fig. 3). We also show solutions at *γ* = 2.2 and *γ* = 1.9 in the supplement (Fig. S3, S4).

### Neurosynth

Probabilistic measures of the association between voxels and cognitive processes were obtained from Neurosynth, a meta-analytic tool that synthesizes results from more than 14 000 published fMRI studies by searching for high-frequency key words (such as “pain” and “attention”) that are published alongside fMRI voxel coordinates (https://github.com/neurosynth/neurosynth, using the volumetric association test maps [126]). This measure of association is the probability that a given cognitive process is reported in the study if there is activation observed at a given voxel. Although more than a thousand cognitive processes are reported in Neurosynth, we focus primarily on cognitive function and therefore limit the terms of interest to cognitive and behavioural terms. These terms were selected from the Cognitive Atlas, a public ontology of cognitive science [86], which includes a comprehensive list of neurocognitive processes. We used 123 terms, ranging from umbrella terms (“attention”, “emotion”) to specific c ognitive p rocesses (“visual a ttention”, “episodic memory”), behaviours (“eating”, “sleep”), and emotional states (“fear”, “anxiety”). The coordinates reported by Neurosynth were parcellated according to the Schaefer atlas and z-scored [96]. The full list of cognitive processes is shown in Supplementary Table S2.

### Neurotransmitter receptors & transporters

PET-derived receptor density data were collated by Hansen et al. [51] and downloaded from neuromaps (https://github.com/netneurolab/neuromaps [65]) for 18 neurotransmitter receptors and transporters across 9 neurotransmitter systems,. These include dopamine (D_2_ [92], DAT [95]), norepinephrine (NET [32]), serotonin (5-HT_1A_ [13], 5-HT_1B_ [41], 5-HT_2A_ [13], 5-HT_4_ [13], 5-HT_6_ [87, 88], 5-HTT [13]), acetylcholine (*α*_4_*β*_2_ [56], M_1_ [73], VAChT [1]), glutamate (mGluR_5_ [34]), GABA (GABA_A_ [77]), histamine (H_3_ [42]), cannabinoid (CB_1_ [78]), and opioid (MOR [116]). Methodological details about each tracer can be found in Table S1. Volumetric PET images were parcellated according to both the Schaefer atlas as well as the Brainstem Navigator atlas [16, 96].

### Dominance analysis

Dominance analysis seeks to determine the relative contribution (“dominance”) of each independent variable to the overall fit (adjusted *R*^2^) of the multiple linear regression model (https://github.com/ dominance-analysis/dominance-analysis [4]). This is done by fitting the same regression model on every combination of input variables (2^*p*^ − 1 submodels for a model with *p* input variables). Total dominance is defined as the average of the relative increase in *R*^2^ when adding a single input variable of interest to a submodel, across all 2^*p*^ − 1 submodels. The sum of the dominance of all input variables is equal to the total adjusted *R*^2^ of the complete model, making total dominance an intuitive method that partitions the total effect size across predictors. Therefore, unlike other methods of assessing predictor importance, such as methods based on regression coefficients or univariate correlations, dominance analysis accounts for predictor-predictor interactions and is interpretable. Dominance was then normalized by the total fit (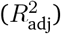) of the model, to make dominance fully comparable both within and across models. The normalized total dominance (percent contribution) is plotted in the heatmap in Fig. 4.

### Spatial null model

Spatial autocorrelation-preserving permutation tests were used to assess statistical significance of associations across brain regions, termed “spin tests” [2, 66, 120]. We created a surface-based representation of the parcellation on the FreeSurfer fsaverage surface, via files from the Connectome Mapper toolkit (https://github. com/LTS5/cmp). We used the spherical projection of the fsaverage surface to define spatial coordinates for each parcel by selecting the coordinates of the vertex closest to the center of the mass of each parcel. These parcel coordinates were then randomly rotated, and original parcels were reassigned the value of the closest rotated parcel according to the Hungarian algorithm (10 000 repetitions) [61]. The procedure was performed at the parcel resolution rather than the vertex resolution to avoid upsampling the data, and to each hemisphere separately.

## Acknowledgments

We thank Vincent Bazinet, Eric G. Ceballos, Asa Farahani, Zhen-Qi Liu, Andrea Luppi, Filip Milisav, and Moohebat Pourmajidian for their comments and suggestions on the manuscript. We also thank Dr. M. Hansen for inspiring the choice of colourmap by way of his particular taste in coat hanger colours. BM acknowledges support from the Natural Sciences and Engineering Research Council of Canada (NSERC), Canadian Institutes of Health Research (CIHR), Brain Canada Foundation Future Leaders Fund, the Canada Research Chairs Program, the Michael J. Fox Foundation, and the Healthy Brains for Healthy Lives initiative. JYH acknowledges support from the Helmholtz International BigBrain Analytics & Learning Laboratory, the Natural Sciences and Engineering Research Council of Canada, and The Neuro Irv and Helga Cooper Foundation. MB acknowledges support from the National Institute of Aging, NIH: R01-AG063982 and the Michael J. Fox Foundation: MJFF-022672 Award. The funders had no role in study design, data collection and analysis, decision to publish or preparation of the manuscript.

---

**Figure S1.**
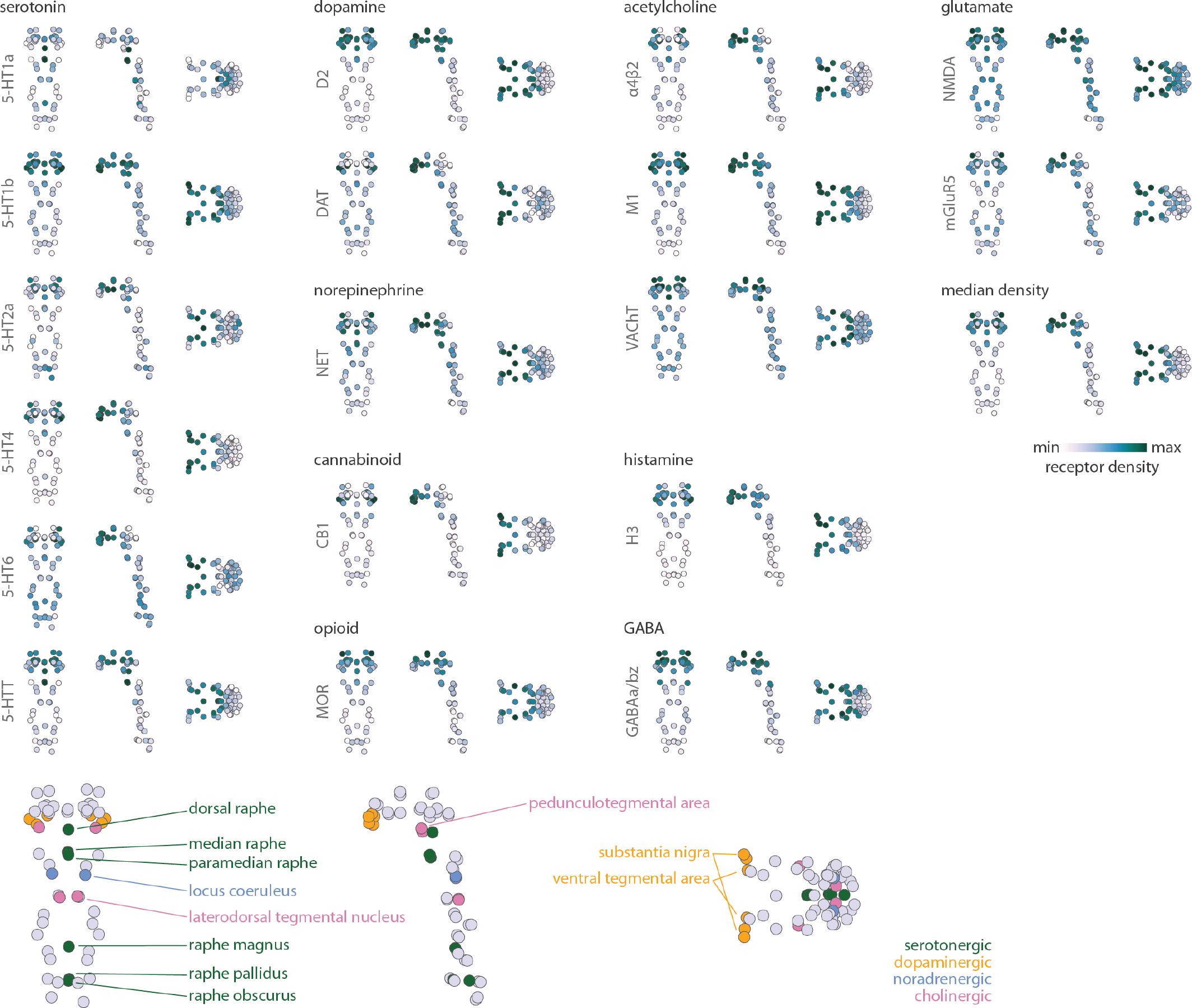
Neurotransmitter receptor and transporter densities in the brainstem. | 18 PET-derived neurotransmitter receptor and transporter density profiles are shown in the brainstem, as well as the median density across all 18 maps. Coronal (posterior view), sagittal, and axial perspectives of brainstem nuclei are shown. A legend of neuromodulatory nuclei in the brainstem is shown in the bottom row.

**Figure S2.**
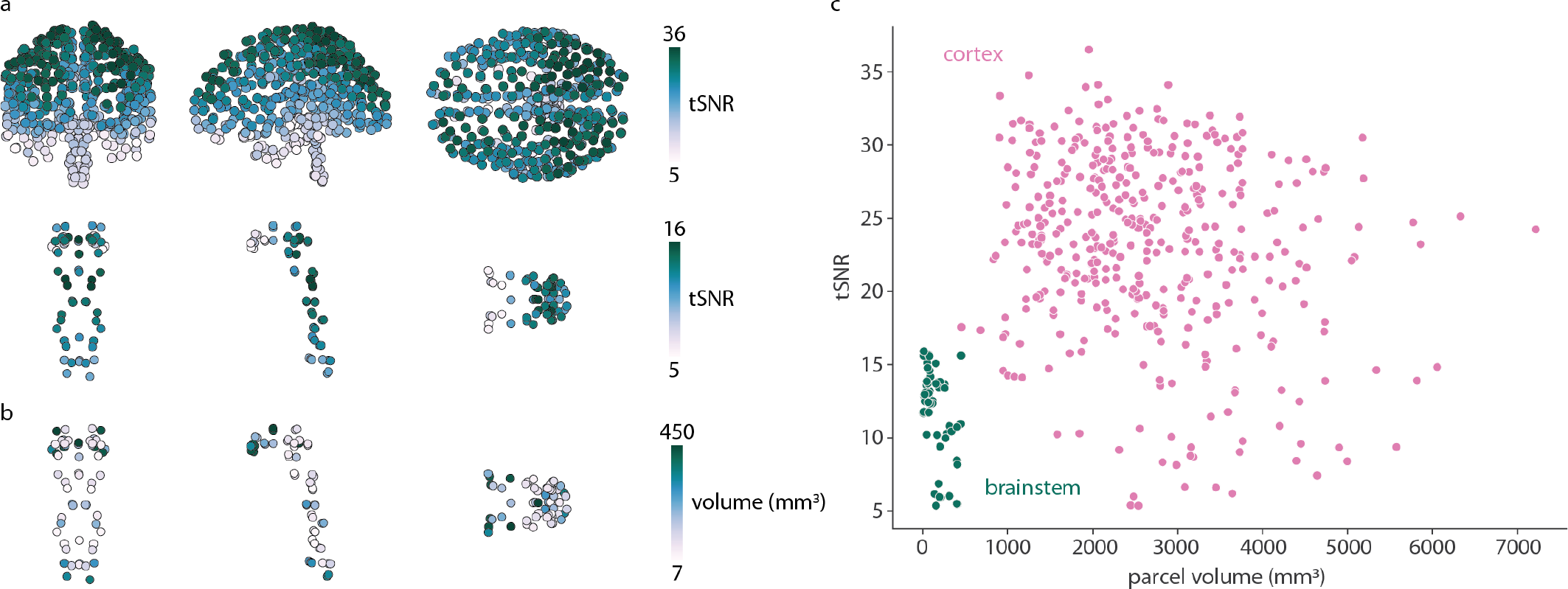
Temporal signal-to-noise ratio and parcel size. (a) Temporal signal-to-noise ratio (tSNR) is calculated as the ratio of the mean of a region’s time-series to its standard deviation (prior to demeaning the time-series in the preprocessing pipeline). tSNR is shown for the cortex and brainstem together (top) and the brainstem only (bottom). Cortical tSNR ∈ [5.37, 36.51], brainstem tSNR ∈ [5.38, 15.92]. (b) Parcel volume in mm^3^ is shown for each brainstem nucleus. (c) Scatter plot showing the relationship between parcel volume and tSNR of brainstem (green; *r* = −0.45, *p* = 0.0004) and cortical (pink; *r* = −0.15, *p* = 0.003) regions.

**TABLE S1.** Neurotransmitter receptors and transporters. | BP_ND_ = non-displaceable binding potential; V_T_ = tracer distribution volume; B_max_ = density (pmol/ml) converted from binding potential (5-HT) or distributional volume (GABA) using autoradiography-derived densities; SUVR = standard uptake value ratio. Values in parentheses (under *N*) indicate number of males.

| Receptor/<br>transporter | Neurotransmitter | Tracer | Measure | $N$ | References |
| --- | --- | --- | --- | --- | --- |
| D<sub>2</sub> | dopamine | [<sup>11</sup>C]FLB-457 | BP<sub>ND</sub> | 55 (26) | [92, 93, 107, 108, 128] |
| DAT | dopamine | [<sup>123</sup>I]-FP-CIT | SUVR | 174 (109) | [35] |
| NET | norepinephrine | [<sup>11</sup>C]MRB | BP<sub>ND</sub> | 77 (50) | [12, 29, 32, 91] |
| 5-HT<sub>1A</sub> | serotonin | [<sup>11</sup>C]CUMI-101 | BP<sub>ND</sub> | 8 (3) | [13] |
| 5-HT<sub>1B</sub> | serotonin | [<sup>11</sup>C]P943 | BP<sub>ND</sub> | 23 (15) | [7, 41, 67, 71, 72, 84, 94] |
| 5-HT<sub>2A</sub> | serotonin | [<sup>11</sup>C]Cimbi-36 | B<sub>max</sub> | 29 (15) | [13] |
| 5-HT<sub>4</sub> | serotonin | [<sup>11</sup>C]SB207145 | B<sub>max</sub> | 59 (41) | [13] |
| 5-HT<sub>6</sub> | serotonin | [<sup>11</sup>C]GSK215083 | BP<sub>ND</sub> | 30 (30) | [87, 88] |
| 5-HTT | serotonin | [<sup>11</sup>C]DASB | B<sub>max</sub> | 100 (29) | [13] |
| $\alpha_4\beta_2$ | acetylcholine | [<sup>18</sup>F]flubatine | V<sub>T</sub> | 30 (20) | [6, 56] |
| M<sub>1</sub> | acetylcholine | [<sup>11</sup>C]LSN3172176 | BP<sub>ND</sub> | 24 (13) | [73] |
| VACHT | acetylcholine | [<sup>18</sup>F]FEOBV | SUVR | 18 (5) | [1] |
| mGluR<sub>5</sub> | glutamate | [<sup>11</sup>C]ABP688 | BP<sub>ND</sub> | 28 (15) | [34] |
| GABA<sub>A/BZ</sub> | GABA | [<sup>11</sup>C]flumazenil | B<sub>max</sub> | 16 (7) | [77] |
| H<sub>3</sub> | histamine | [<sup>11</sup>C]GSK189254 | V<sub>T</sub> | 8 (7) | [42] |
| CB<sub>1</sub> | cannabinoid | [<sup>11</sup>C]OMAR | V<sub>T</sub> | 77 (49) | [36, 75, 78, 89] |
| MOR | opioid | [<sup>11</sup>C]carfentanil | BP<sub>ND</sub> | 39 (19) | [116] |

**Figure S3.**
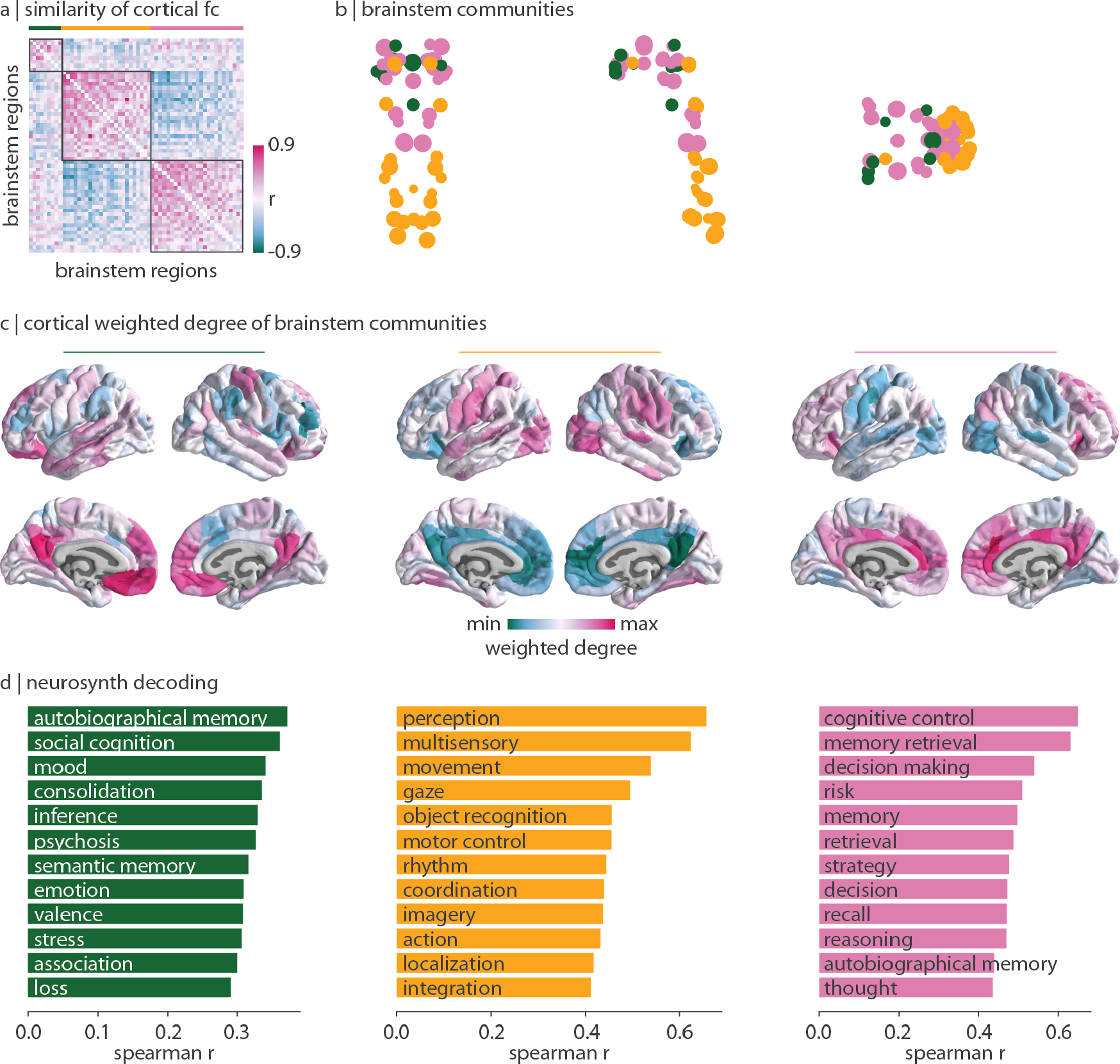
Brainstem communities when. *γ* = 1.9 | The Louvain community detection algorithm was repeated for *γ* = 1.9 which identified three stable communities.

**Figure S4.**
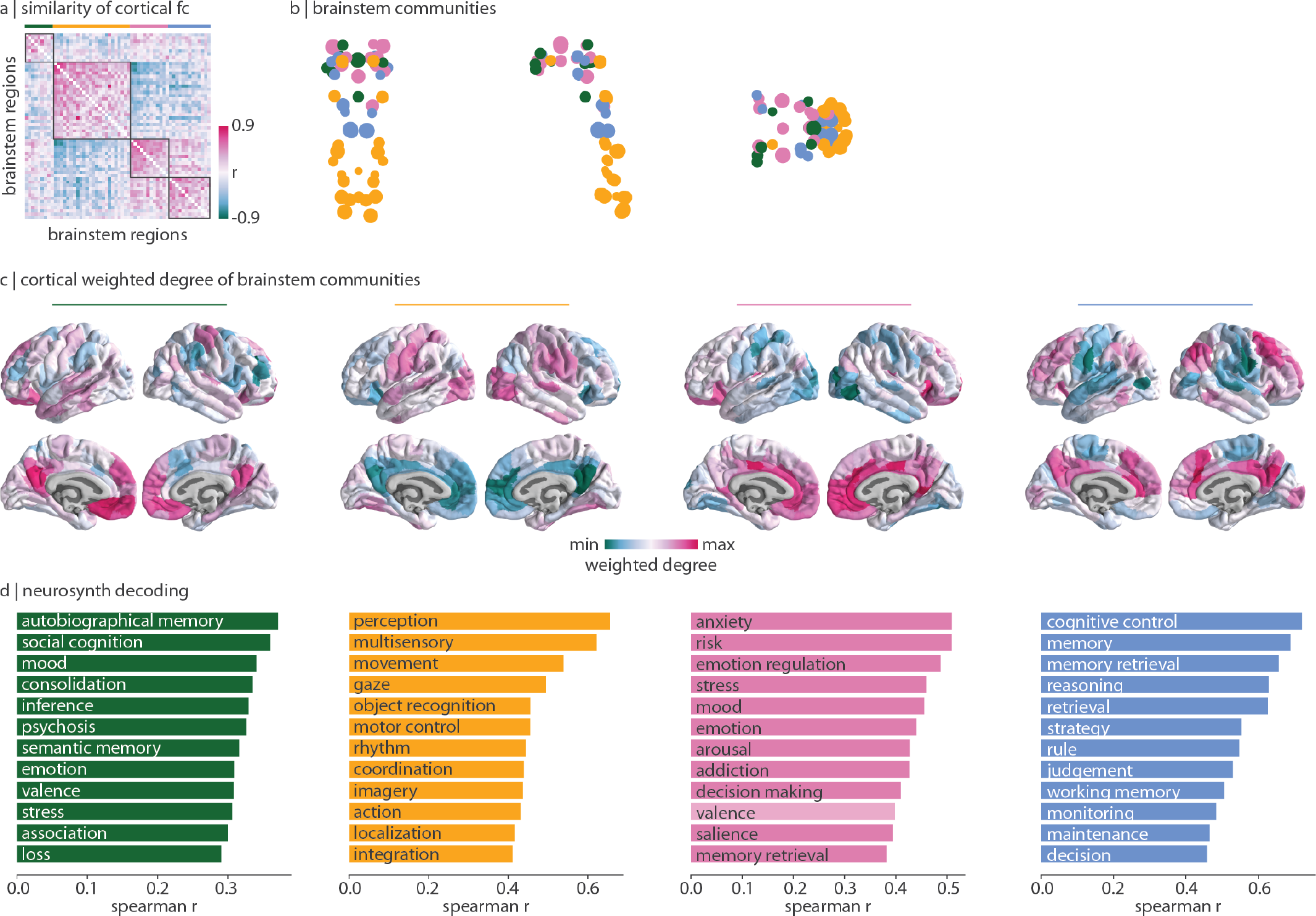
Brainstem communities when. *γ* = 2.2 | The Louvain community detection algorithm was repeated for *γ* = 2.2 which identified four stable communities.

**Figure S5.**
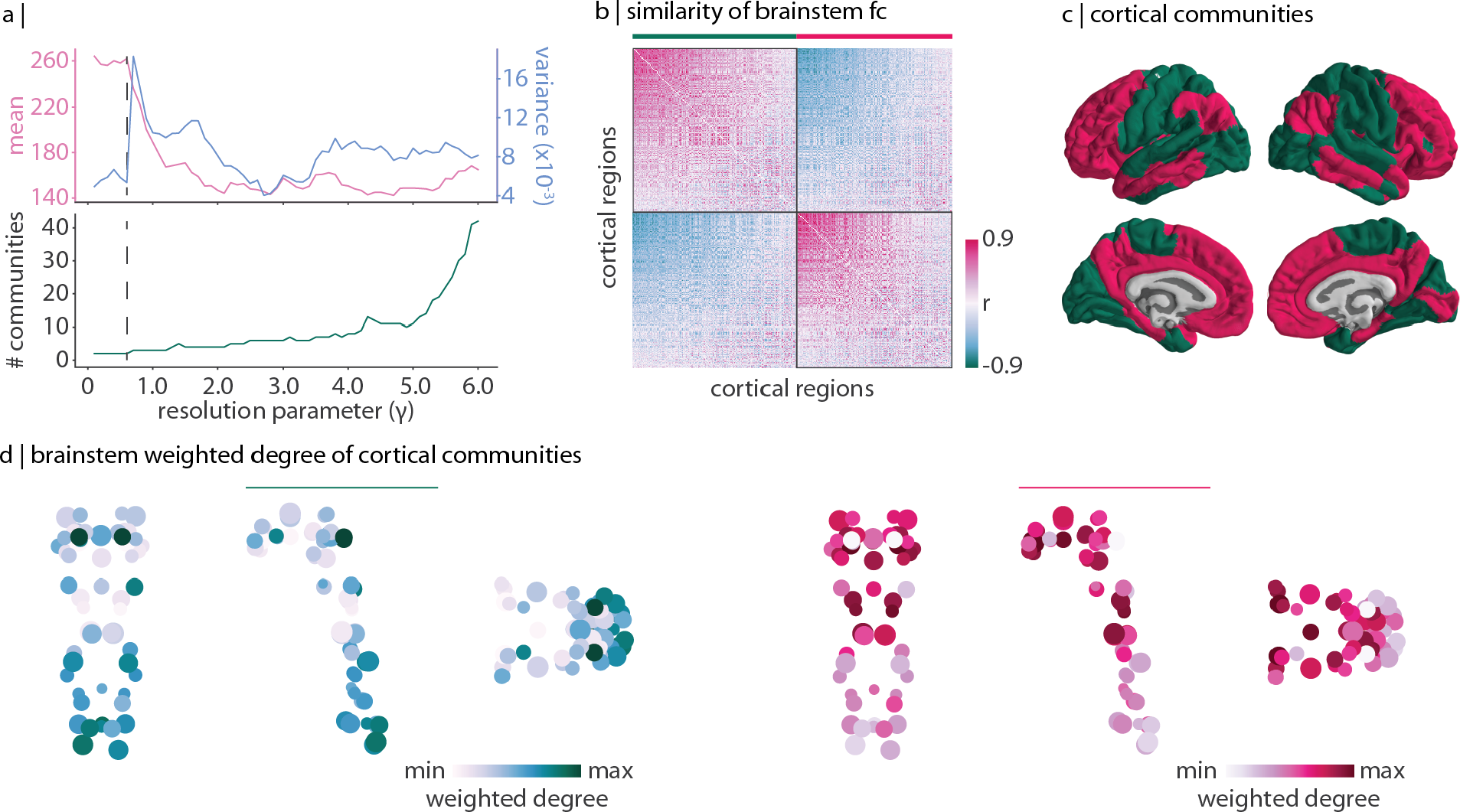
Cortical communities of brainstem functional connectivity. | The Louvain community detection algorithm was applied to a correlation matrix representing how similarly (Spearman’s *r*) two cortical regions are functionally connected with the brainstem above and beyond the dominant pattern of brainstem connectivity. (a) Top: mean and variance of the z-scored rand index across 250 repetitions of the Louvain algorithm at each resolution parameter *γ* ∈ [0.1, 6.0]. Bottom: number of communities identified for each *γ*. The dashed vertical line exists at *γ* = 0.6. (b) Cortical region × region correlation matrix representing how similarly cortical regions are functionally connected with the brainstem. Regions are ordered according to the two communities identified at *γ* = 0.6. (c) Community affiliations for each cortical region. (d) Brainstem weighted degree of the green (left, unimodal) community and the red (right, transmodal) community. Specifically, for each brainstem nucleus, we sum its regressed functional connectivity with all cortical regions in the green/red community.

**Figure S6.**
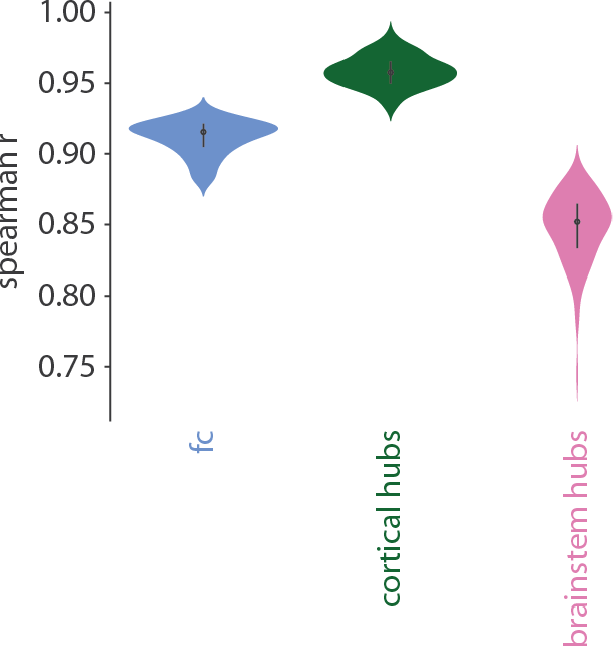
Split-half analysis. | The 20 participants included in the present study were randomly divided into two groups of 10 (100 repetitions). Group-average functional connectivity, cortex-to-brainstem weighted degree patterns, and brainstem-to-cortex weighted degree patterns were recalculated within these groups are correlated.

**Figure S7.**
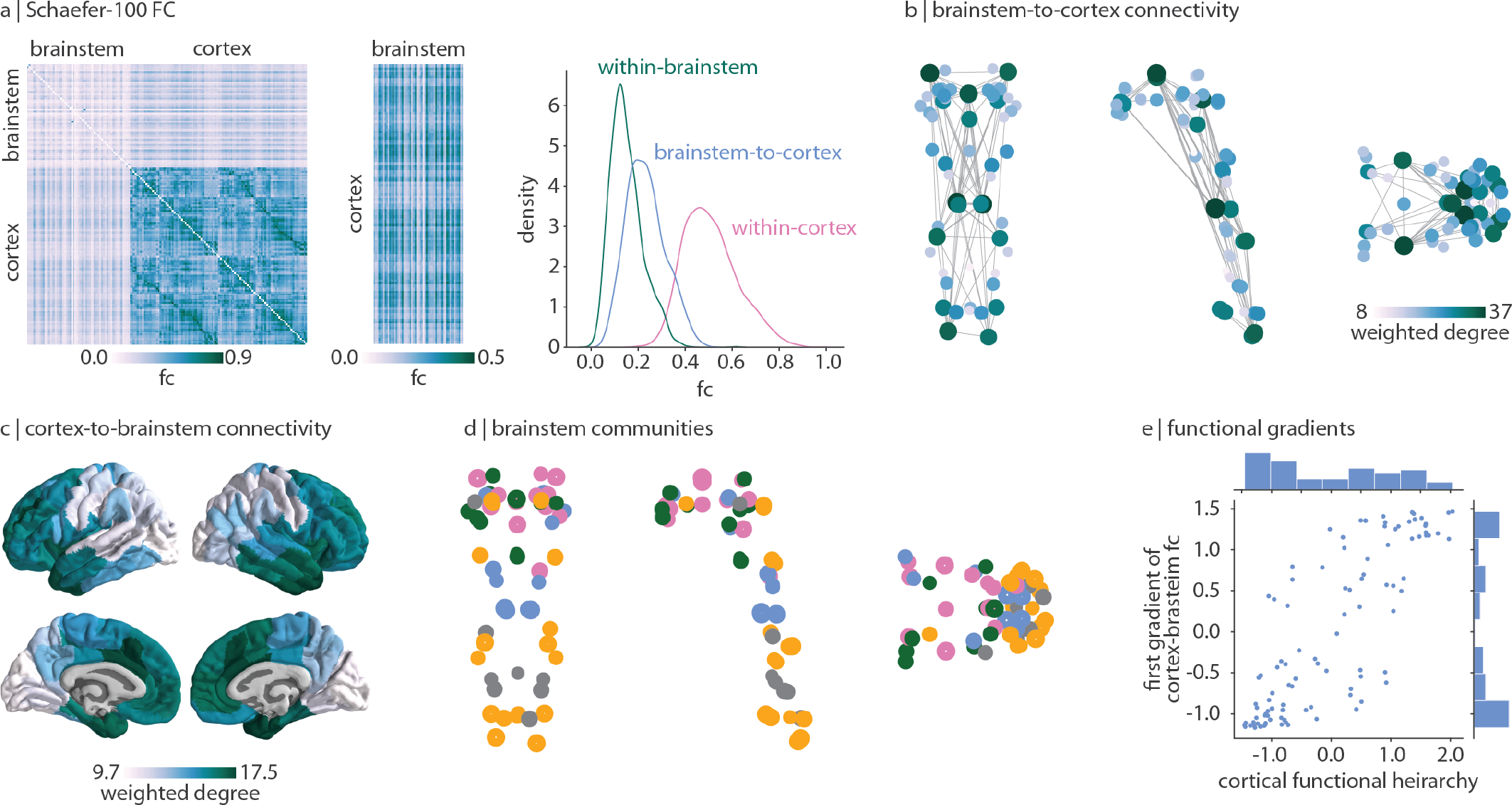
Replication using 100 cortical regions. | Analyses were repeated using the 100-region Schaefer parcellation [96]. (a) Functional connectivity and functional connectivity density distributions. (b) Brainstem-to-cortex weighted degree. (c) Cortexto-brainstem weighted degree. (d) Community affiliations of brainstem nodes under identical parameters as shown in Fig. 3. (e) Correlation between the first gradient of cortex-to-brainstem functional connectivity and the cortical functional hierarchy.

**Figure S8.**
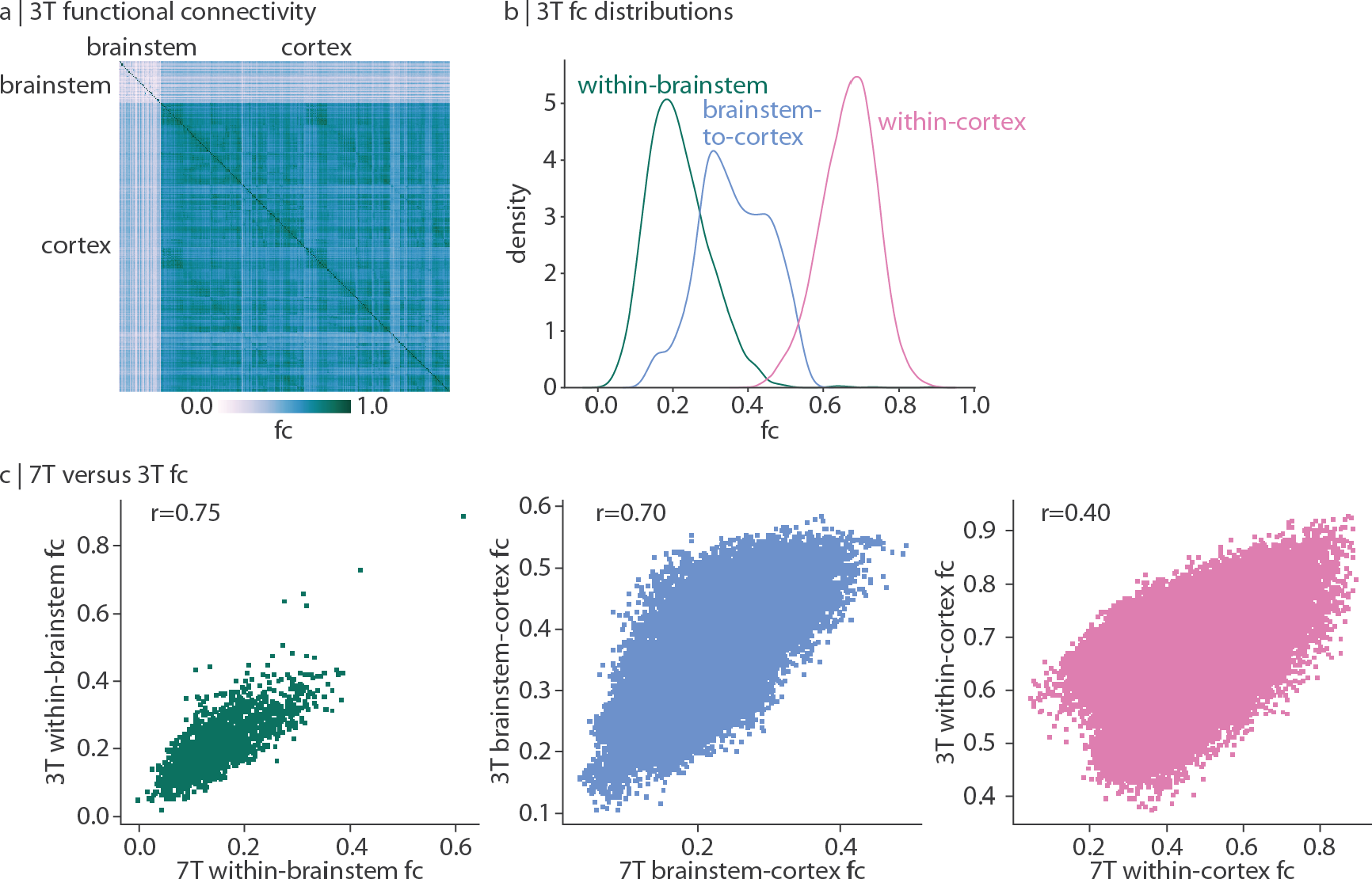
Replication using 3 Tesla fMRI data. | Analyses were repeated using 3 Tesla fMRI data acquired in the same 20 subjects, under the Schaefer-400 parcellation [96]. (a) Functional connectivity. (b) Functional connectivity density distributions. (c) Spearman correlations between 7 Tesla functional connectivity data used in the main analyses and 3 Tesla functional connectivity replication data.

**Figure S9.**
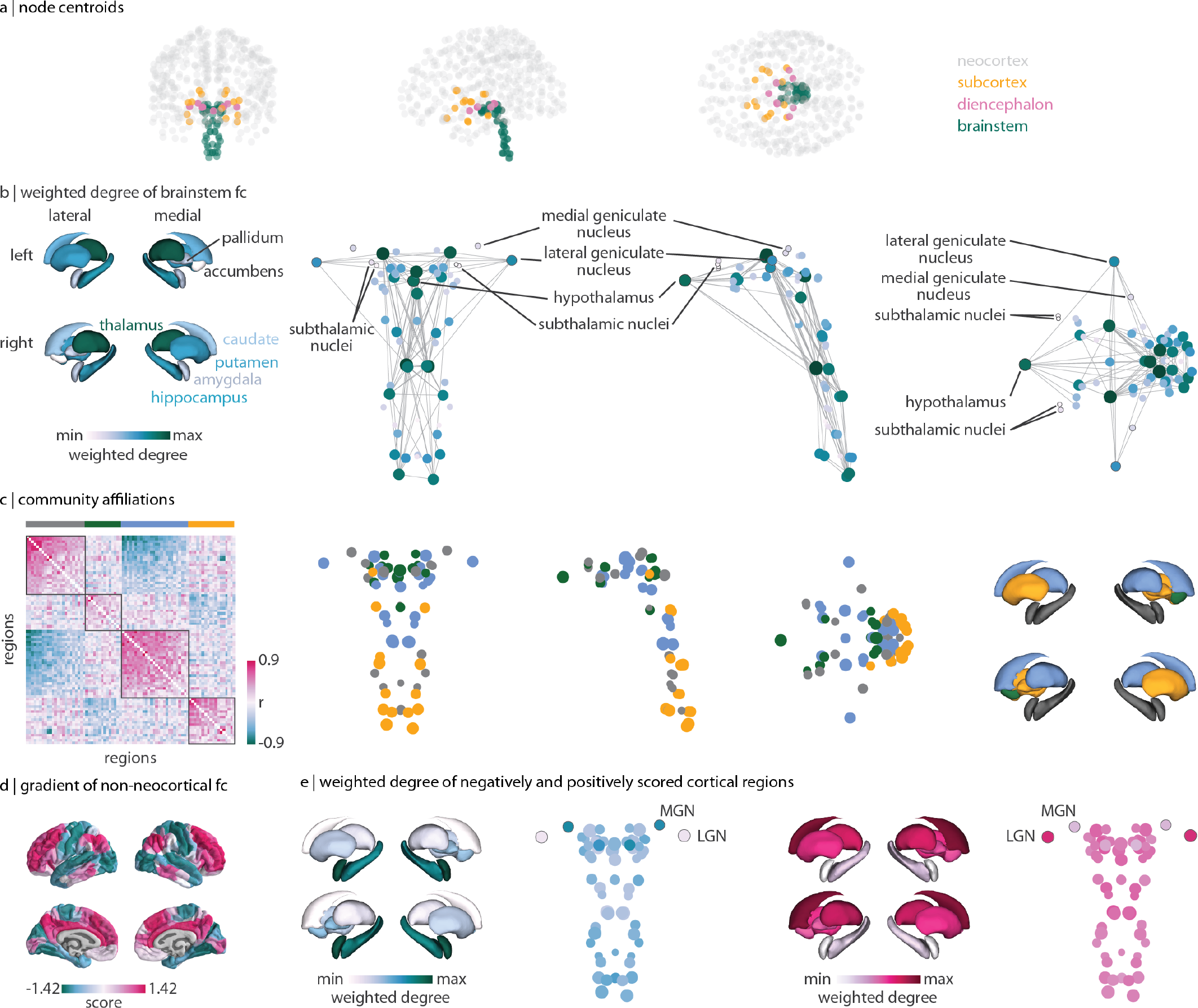
Extending analyses to subcortical and diencephalic structures. | Functional images were also acquired for the 14 bilateral FreeSurfer subcortical structures (caudate, putamen, pallidum, nucleus accumbens, thalamus, amygdala, hippocampus (not technically subcortex but allocortex)) as well as 8 bilateral diencephalic structures from the Brainstem Navigator (lateral geniculate nucleus, medial geniculate nucleus, subthalamic nuclei subregions 1 & 2). (a) For each brain regions, the centroid coordinate is plotted with colours indicating structure. Grey: 400 neocortical structures; yellow: 14 FreeSurfer subcortical structures; pink: 8 Brainstem Navigator diencephalic regions; green: 58 Brainstem Navigator brainstem nuclei. (b) Left: FreeSurfer subcortical plot of weighted degree of brainstem functional connectivity, representing how much each FreeSurfer subcortical parcel is connected with the brainstem. Each structure is labeled. Right: Brainstem Navigator brainstem and diencephalic centroid coordinates coloured according to their weighted degree of brainstem functional connectivity, representing how much each nucleus is connected with the 58 brainstem structures. The 8 diencephalic nuclei are labeled. (c) Left: region × region similarity matrix representing how similarly two non-neocortical (i.e. brainstem, subcortical, or diencephalic) regions are functionally connected with the cortex. Outlines are placed around the identified communities. Middle: point brain plot of community assignments for brainstem and diencephalic nuclei. Right: FreeSurfer surface plot of community assignments for FreeSurfer subcortical regions. (d) Left: region × region similarity matrix representing how similarly two cortical regions are functionally connected with non-neocortical regions (i.e. brainstem, subcortical, diencephalic). Outlines are shown around the seven Yeo-Krienen resting-state networks (order: control, default mode, dorsal attention, limbic, ventral attention, somato-motor, visual). Middle: coritcal surface plot of the first gradient from diffusion map embedding of how similarly cortical regions are functionally connected with non-neocortical regions. Right: FreeSurfer subcortical weighted degree patterns, calculated as the sum of a FreeSurfer subcortical region’s functional connectivity with all negatively- (left) or positively- (right) scored regions of the cortical gradient shown on the left. FreeSurfer subcortical structures were plotted using the enigmatoolbox [62].

**Figure S10.**
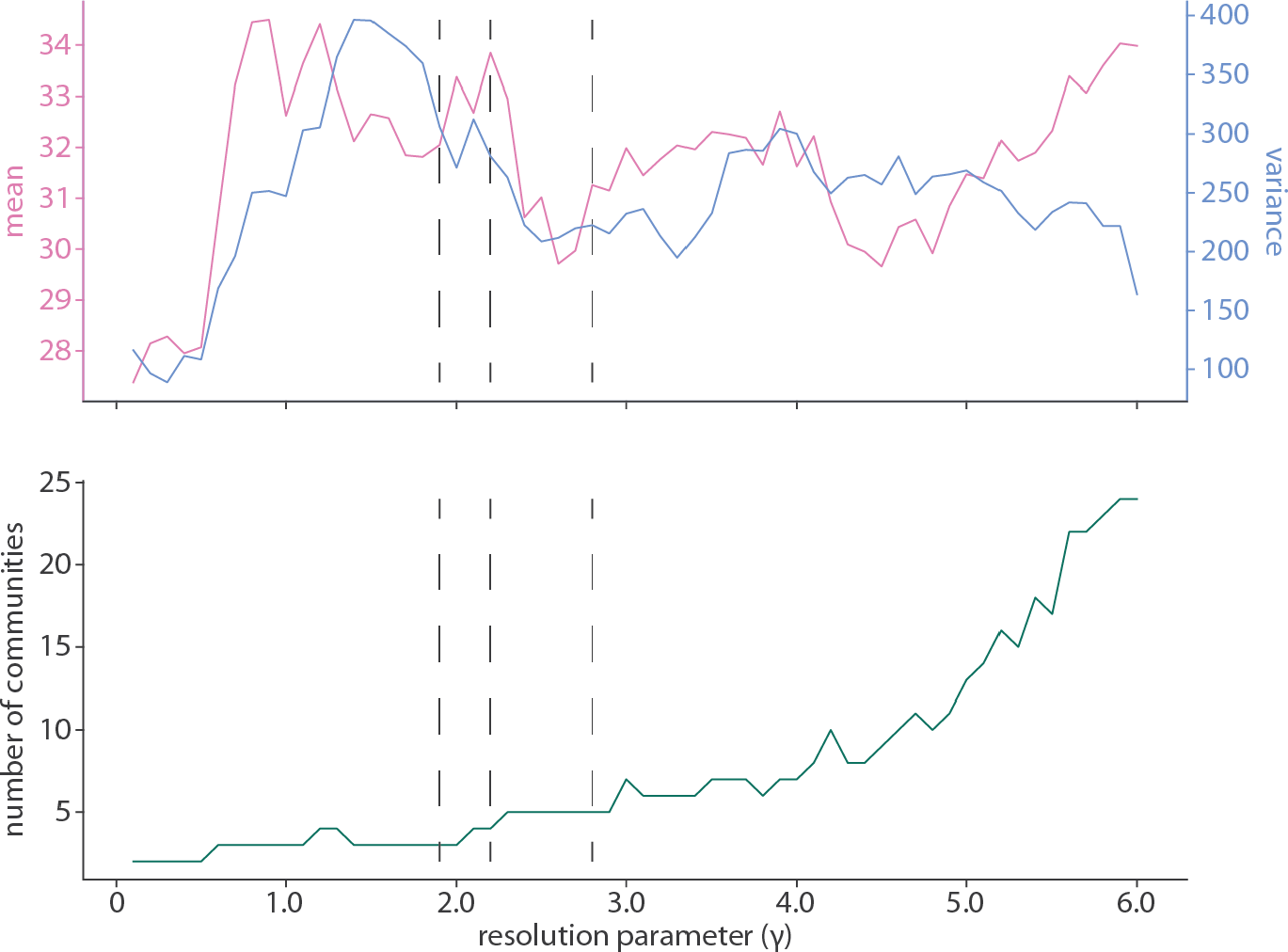
Community detection performance across resolution parameter. *γ* | Top: mean and variance of the z-scored rand index across 250 repetitions of the Louvain community detection algorithm at each *γ* (for *γ* ∈ [0.1, 6.0]). Community detection solutions are considered better quality (i.e. more stable) when the mean of the z-scored rand index is high and the variance is low. Dashed vertical lines are placed at values of *γ* where the community detection solution is shown in the text (*γ* = 1.9 shown in Fig. S3, *γ* = 2.2 shown in Fig. S4, and *γ* = 2.8 shown in Fig. 3). Bottom: the number of communities identified by the algorithm across values of *γ*.

**TABLE S2.**
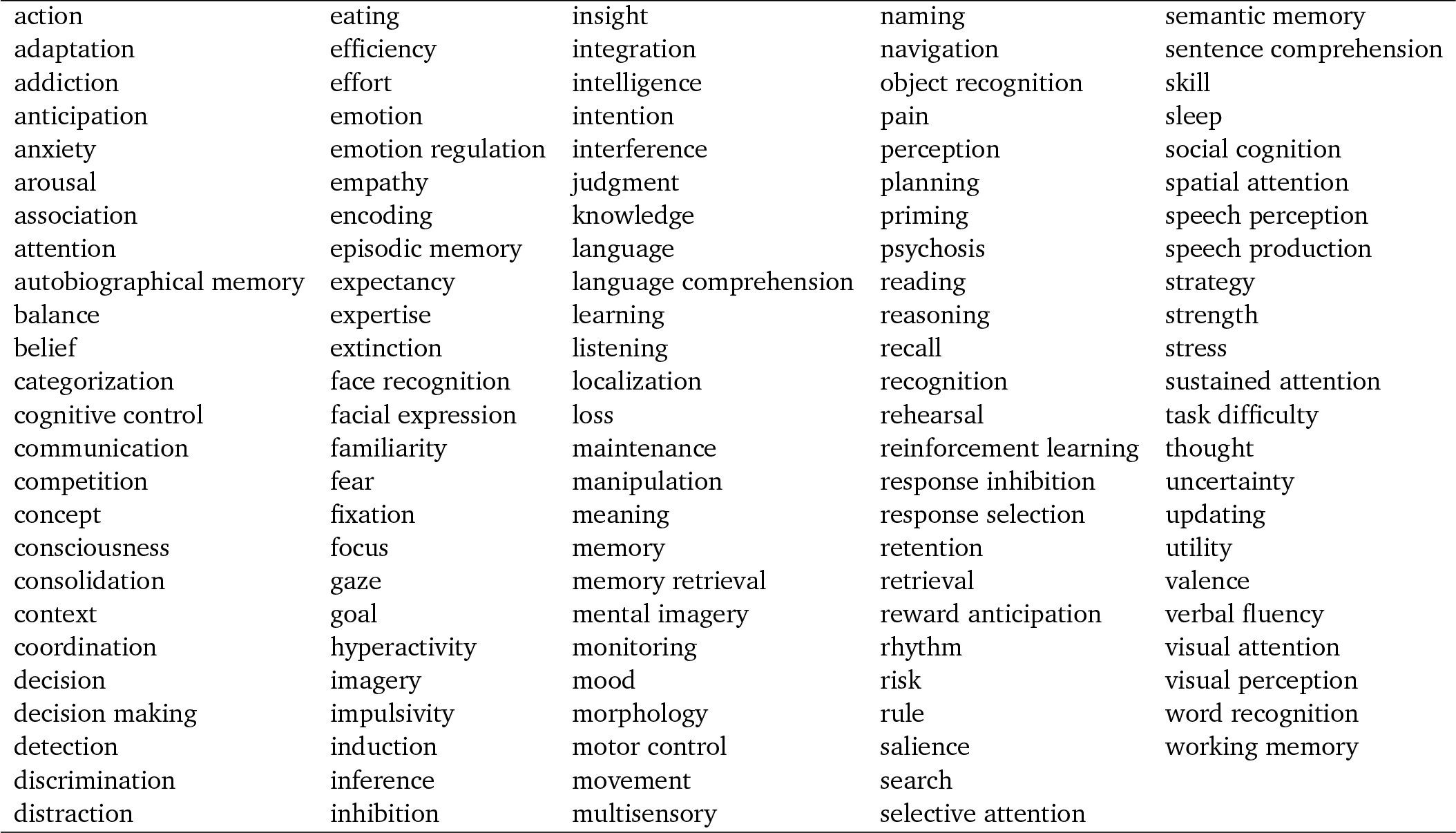
Neurosynth terms. | Terms that overlapped between the Neurosynth database [126] and the Cognitive Atlas [86] were used in the cognitive decoding analysis.

|  |  |  |  |  |
| --- | --- | --- | --- | --- |
| action | eating | insight | naming | semantic memory |
| adaptation | efficiency | integration | navigation | sentence comprehension |
| addiction | effort | intelligence | object recognition | skill |
| anticipation | emotion | intention | pain | sleep |
| anxiety | emotion regulation | interference | perception | social cognition |
| arousal | empathy | judgment | planning | spatial attention |
| association | encoding | knowledge | priming | speech perception |
| attention | episodic memory | language | psychosis | speech production |
| autobiographical memory | expectancy | language comprehension | reading | strategy |
| balance | expertise | learning | reasoning | strength |
| belief | extinction | listening | recall | stress |
| categorization | face recognition | localization | recognition | sustained attention |
| cognitive control | facial expression | loss | rehearsal | task difficulty |
| communication | familiarity | maintenance | reinforcement learning | thought |
| competition | fear | manipulation | response inhibition | uncertainty |
| concept | fixation | meaning | response selection | updating |
| consciousness | focus | memory | retention | utility |
| consolidation | gaze | memory retrieval | retrieval | valence |
| context | goal | mental imagery | reward anticipation | verbal fluency |
| coordination | hyperactivity | monitoring | rhythm | visual attention |
| decision | imagery | mood | risk | visual perception |
| decision making | impulsivity | morphology | rule | word recognition |
| detection | induction | motor control | salience | working memory |
| discrimination | inference | movement | search |  |
| distraction | inhibition | multisensory | selective attention |  |

